# Cryo-EM Structures Delineate a pH-Dependent Switch that Mediates Endosomal Positioning of SARS-CoV-2 Spike Receptor-Binding Domains

**DOI:** 10.1101/2020.07.04.187989

**Authors:** Tongqing Zhou, Yaroslav Tsybovsky, Adam S. Olia, Jason Gorman, Micah Rapp, Gabriele Cerutti, Gwo-Yu Chuang, Phinikoula S. Katsamba, Alexandra Nazzari, Jared M. Sampson, Arne Schön, Pengfei Wang, Jude Bimela, Wei Shi, I-Ting Teng, Baoshan Zhang, Jeffrey C. Boyington, Mallika Sastry, Tyler Stephens, Jonathan Stuckey, Shuishu Wang, Richard A. Friesner, David D. Ho, John R. Mascola, Lawrence Shapiro, Peter D. Kwong

## Abstract

The SARS-CoV-2 spike employs mobile receptor-binding domains (RBDs) to engage the ACE2 receptor and to facilitate virus entry. Antibodies can engage RBD but some, such as CR3022, fail to inhibit entry despite nanomolar spike affinity. Here we show the SARS-CoV-2 spike to have low unfolding enthalpy at serological pH and up to 10-times more unfolding enthalpy at endosomal pH, where we observe significantly reduced CR3022 affinity. Cryo-EM structures –at serological and endosomal pH– delineated spike recognition of up to three ACE2 molecules, revealing RBD to freely adopt the ‘up’ conformation. In the absence of ACE2, single-RBD-up conformations dominated at pH 5.5, resolving into a locked all-down conformation at lower pH. Notably, a pH-dependent refolding region (residues 824-858) at the spike-interdomain interface displayed dramatic structural rearrangements and mediated RBD positioning and spike shedding of antibodies like CR3022. An endosomal mechanism involving spike-conformational change can thus facilitate immune evasion from RBD-‘up’-recognizing antibody.

**Highlights:**

- Reveal spike at serological pH to have only ~10% the unfolding enthalpy of a typical globular protein, explaining how antibodies like CR3022 can bind with avidity
- Define an endosomal mechanism whereby spike binds ACE2, but sheds CR3022, enabling immune evasion from potentially neutralizing antibody
- Determine cryo-EM structures of the SARS-CoV-2 spike along its endosomal entry pathway-at pH 5.5, 4.5, and 4.0, and in complexes with ACE2 receptor at pH 7.4 and 5.5
- Show spike to exclusively adopt an all RBD-down conformation at the low pH of the late endosome-early lysosome
- Reveal structural basis by which a switch domain mediates RBD position in response to pH

## Introduction

The SARS-CoV-2 spike is a type 1 fusion machine, responsible for virus-cell entry via ACE2-receptor interactions (Lan et al., 2020; Shang et al., 2020b; Wang et al., 2020; Zhou et al., 2020a). Entry occurs both endosomally and at the cell surface, with inhibition of the endosomal cathepsin L and the cell-surface TMPRSS2 required to fully inhibit entry (Hoffmann et al., 2020; Ou et al., 2020); cleavage of the spike by furin can also occur, but furin cleavage does not appear to be essential for entry and occurs distal from the fusion peptide. Cryo-EM structures reveal two prevalent conformations for uncleaved and furin-cleaved SARS-CoV-2 spikes (Walls et al., 2020; Wrapp et al., 2020; Wrobel et al., 2020): a single-up conformation and an all-down conformation, related to the positioning of the receptor-binding domains (RBDs). The ‘up’ positioning of RBD is required for interaction with ACE2 receptor and is also related to the epitope availability of RBD-directed antibodies.

Potent neutralizing antibodies have been identified that target RBD, and in some cases their structures with spike or RBD have been determined (Barnes et al., 2020; Cao et al., 2020; Hansen et al., 2020; Ju et al., 2020; Liu et al., 2020; Shi et al., 2020; Walls et al., 2019; Wu et al., 2020). Other RBD-directed antibodies, such as antibody CR3022 (ter Meulen et al., 2006), however, have been shown to bind spike with high affinity, yet fail to inhibit SARS-CoV-2 entry (Yuan et al., 2020). This high affinity for the spike trimer, yet lack of virus neutralization, suggests a spike-based mechanism to evade potentially neutralizing antibody. Reports of antibody CR3022 disassembling spike (Huo et al., 2020), moreover, suggest unusual spike fragility.

As antibody-bound disassembled spikes seemed unlikely to be capable of inducing direct virus entry, we explored endosomal entry and carried out biophysical and structural studies of the SARS-CoV-2 spike along its endosomal entry pathway. We measured the unfolding enthalpy of the spike as well as its binding to CR3022 antibody and to ACE2 receptor as a function of pH. We determined cryo-EM structures of the spike, alone and in complex with ACE2 receptor, at serological and endosomal pH. We delineate the molecular mechanism that mediates positioning of RBDs, highlighting the key role of a refolding region with multiple aspartic acid residues, a pH-dependent switch, which when protonated locks RBDs in the down position. Overall, our findings provide a pH-dependent mechanism of conformational masking, whereby reduced folding at serological pH underlies the ease by which antibodies like CR3022 bind to spike and their prevalent elicitation. ACE2 recognition and endosomal entry, however, result in reduction of pH and induction of antibody shedding through structural rearrangements of the spike mediated by the pH-dependent switch.

## Results

### Spike at serological pH is at a minimum of unfolding enthalpy

To provide insight into the stability of the spike over the course of virus-cell endosomal entry, we used differential scanning calorimetry (DSC) to measure the unfolding enthalpy of the soluble trimeric ectodomain (spike), which included GSAS and PP mutations and the T4 phage fibritin trimerization domain (Wrapp et al., 2020), as a function of pH. Notably, at serological pH (pH 7.4) we observed spike to be at a minimum of unfolding enthalpy (73 kcal/mol) or ~10% the normalized unfolding enthalpy of the average globular protein (Robertson and Murphy, 1997) (Figure 1A, **left**). The amount of folding energy strengthened rapidly as pH dropped, increasing ~10-fold at pH 6 before decreasing as pH reduced further. Analysis of melting curves (Figure 1A, **right**) indicated three distinct peaks: a peak at ~48°C, which dominated under basic conditions and decreased at acidic conditions; a peak at ~65°C, which was barely present at serological pH, but rapidly increased to dominate as pH dropped to 5.5-6.0; and a peak at ~55°C, which first appeared as a shoulder at pH 6, but then increased to dominate and shifted to lower temperature at pH lower than 5.5.

**Figure 1.**
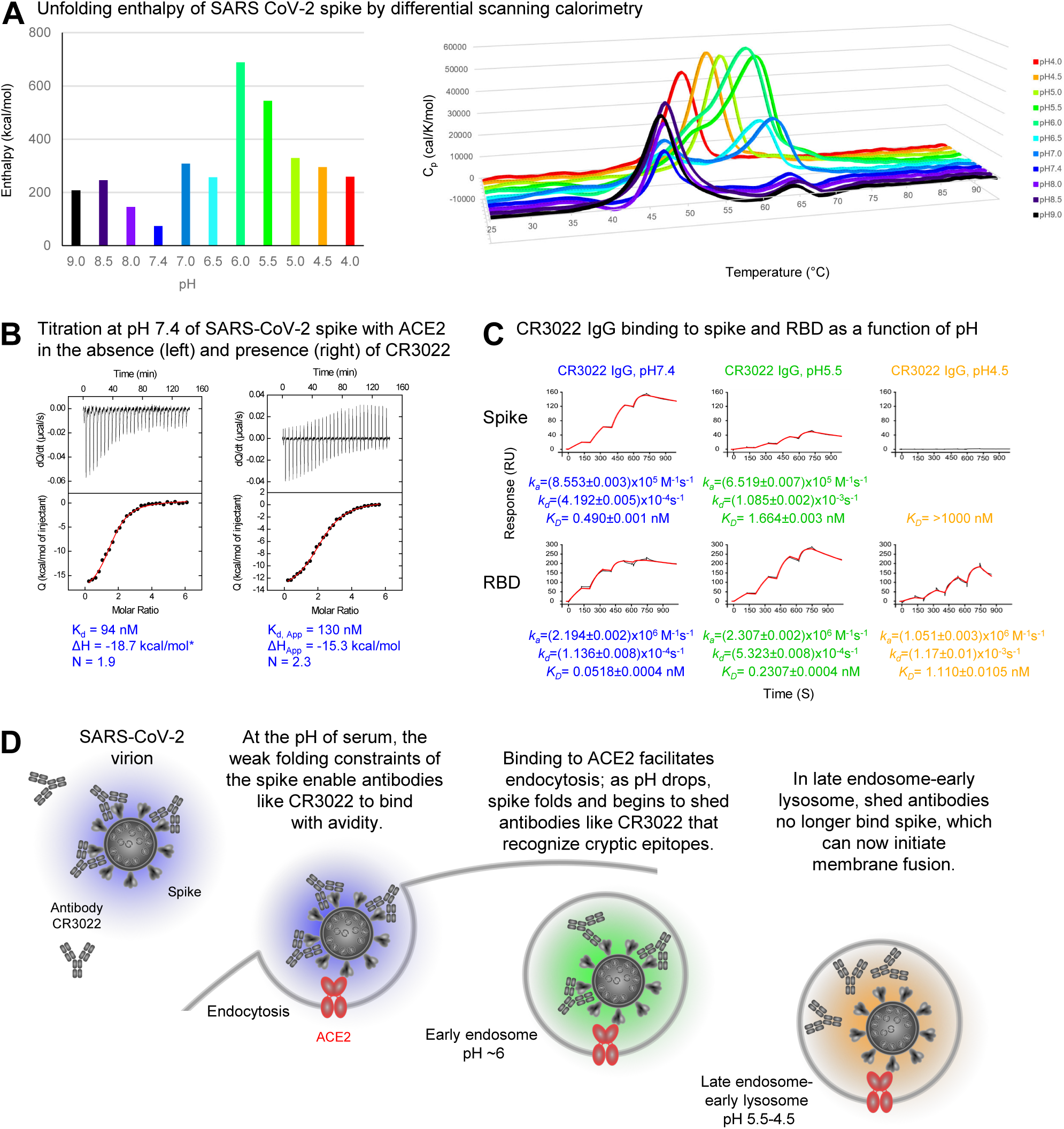
SARS-CoV-2 Spike Is Partially Folded at Serological pH, Where It Binds ACE2 and CR3022, and More Folded Lower pH, Where It Still Binds ACE2, but Not CR3022. (**A**) Unfolding enthalpy of spike as measured by differential scanning calorimetry (DSC). Left, overall unfolding enthalpy measured as area under the curve (AUC) as a function of pH; at pH, the spike showed only ~10% the normalized unfolding enthalpy of the average globular protein. Right, unfolding enthalpy as unction of temperature. (**B**) Isothermal titration calorimetry at pH 7.4 of ACE2 recognizing spike (left) or spike previously titrated with Fab CR3022 (right). (**C**) Apparent affinities of spike (top) and real affinities of RBD (bottom) to CR3022 IgG as a ction of pH as measured by SPR. (**D**) Schematic showing ACE2-dependent endosomal entry of SARS-CoV-2 and the pH-dependent shedding of antibodies like CR3022. See also Figures S1-S3.

In sharp contrast to spike, analysis of the spike N-terminal domain (NTD) and RBD as separate proteins indicated each of them to have typical unfolding enthalpies (>90% that of the average normalized globular protein) (**Figure S1**).

To understand how the reduced folding enthalpy might influence spike structure, we used negative stain-electron microscopy (EM) to visualize the spike as a function of pH (**Figure S2**). At pH 7.4, we observed only a few ordered spikes. However, when pH decreased, we observed the quantity of well-formed trimers to increase substantially; these showed some clustering at pH 5.5-6.0 before resolving into separate particles at pH 4.0-4.5. Overall, we observed concordance between the presence of well-formed trimers and folding enthalpy measured by DSC, with only a few ordered trimers at serological pH and substantially increased well-formed trimers at endosomal pH.

### Binding of ACE2 receptor and CR3022 antibody at serological and endosomal pH

For variation in pH over the course of virus entry to impact the binding of antibody, the antibody would need to allow spike recognition of the ACE2 receptor, thereby enabling the virus to initiate endosomal entry. CR3022 has been shown to not inhibit RBD binding to ACE2 (Yuan et al., 2020), but this has not been shown with spike. We used isothermal titration calorimetry (ITC) to determine whether binding of CR3022 to spike was compatible with ACE2 interaction at serological pH. We chose to use a monomeric version of ACE2 to test more sensitively the impact of antibody inhibition. First, we titrated ACE2 into soluble spike, and observed 1.9 ACE2 molecules to bind per spike trimer, with an affinity of 94 nM (Figure 1B, **left**). Next, we fully titrated the antigen-binding fragment (Fab) of CR3022 into soluble spike (**Figure S3A**) and further titrated ACE2 into the spike-CR3022 complex formed to observe 2.3 ACE2 molecules to bind each spike-CR3022 complex, with an affinity of 130 nM (Figure 1B, **right**). Thus, at serological pH, the SARS-CoV-2 spike appears capable of recognizing ACE2 even in the presence of antibody CR3022, indicating that CR3022-bound spikes could initiate endosomal-based ACE2-dependent entry.

To gain insight into the impact of endosomal pH on ACE2 and CR3022 interactions with spike, we characterized their binding to both spike and RBD, expressed as a separate molecule. For these experiments, we chose to use dimeric ACE2 to more closely mimic native interactions with spike. Endosomes vary in pH from pH ~6 (early endosomes) to pH ~5 (late endosomes), with lysosomal pH as low as ~4 (Benjaminsen et al., 2011; Turk and Turk, 2009). For endosomal pH, we chose to measure pH 5.5 and 4.5. At endosomal pH, surface plasmon resonance (SPR)-determined apparent ACE2 binding affinities to both spike and RBD were somewhat reduced from 0.82 nM at serological pH to 8.4 and 7.0 nM at pH 5.5 and 4.5, respectively, for spike, and from 1.0 nM at serological pH to 2.2 and 15. nM at pH 5.5 and 4.5, respectively, for RBD (**Figure S3B**). With CR3022 IgG, apparent affinities to spike and RBD were sub-nanomolar at serological pH, though with a 10-fold difference (0.49 and 0.052 nM to spike and RBD, respectively) (Figure 1C). At pH 5.5, this 10-fold difference was retained (1.7 and 0.23 nM, respectively). However, at pH 4.5, CR3022 still bound to RBD (1.1 nM), but its apparent affinity to spike was dramatically reduced with a K_D_ >1000 nM – an apparent affinity difference we estimate to be >1000-fold (Figures 1C **and S3C**). Because CR3022 still bound strongly to the isolated RBD, we attribute the dramatically reduced apparent affinity of CR3022 for spike at low pH to conformational constraints of the spike (Figure 1D).

### Cryo-EM structures of SARS-CoV-2 spike with ACE2 at serological and endosomal pH

Structures of ACE2 and CR3022 have been determined in complex with RBD as a separate domain (Lan et al., 2020; Shang et al., 2020b; Wang et al., 2020; Yuan et al., 2020), but less is known about their interactions with trimer. While negative-stain images of soluble ACE2 with spike showed fewer ordered spike complexes at serological pH versus lower pH (**Figure S2**), we judged the spike to be sufficiently ordered to permit residue-level structural analysis at pH 7.4. To provide structural insight into the recognition between ACE2 and spike trimer, we mixed soluble ACE2 with spike trimer at a 6:1 molar ratio at pH 7.4 and collected single-particle cryo-EM data on a Titan Krios. We obtained structures at 3.6-3.9 Å resolution and observed spike to bind ACE2 at stoichiometries of 1:1, 1:2, and 1:3, with prevalences of 15%, 43%, and 38%, respectively (Figure 2A, **Table S1**). While the membrane-proximal region of the spike in these complexes remained 3-fold symmetric, the ACE2-binding regions showed asymmetry with, for example, superposition of the double-ACE2-bound complex onto itself based on membrane-proximal regions leading to displacement of ACE2 molecules by almost 13 Å (Figure 2B). However, we could see no evidence of coordinated movement, with the RBD domain on each protomer appearing to engage ACE2 without significantly impacting the up (or down) positioning of the neighboring protomers. Thus, ACE2-receptor engagement required RBD to be in the ‘up’ position, and this did not appear to destabilize the spike nor to trigger a substantial structural rearrangement beyond raising of RBD.

**Figure 2.**
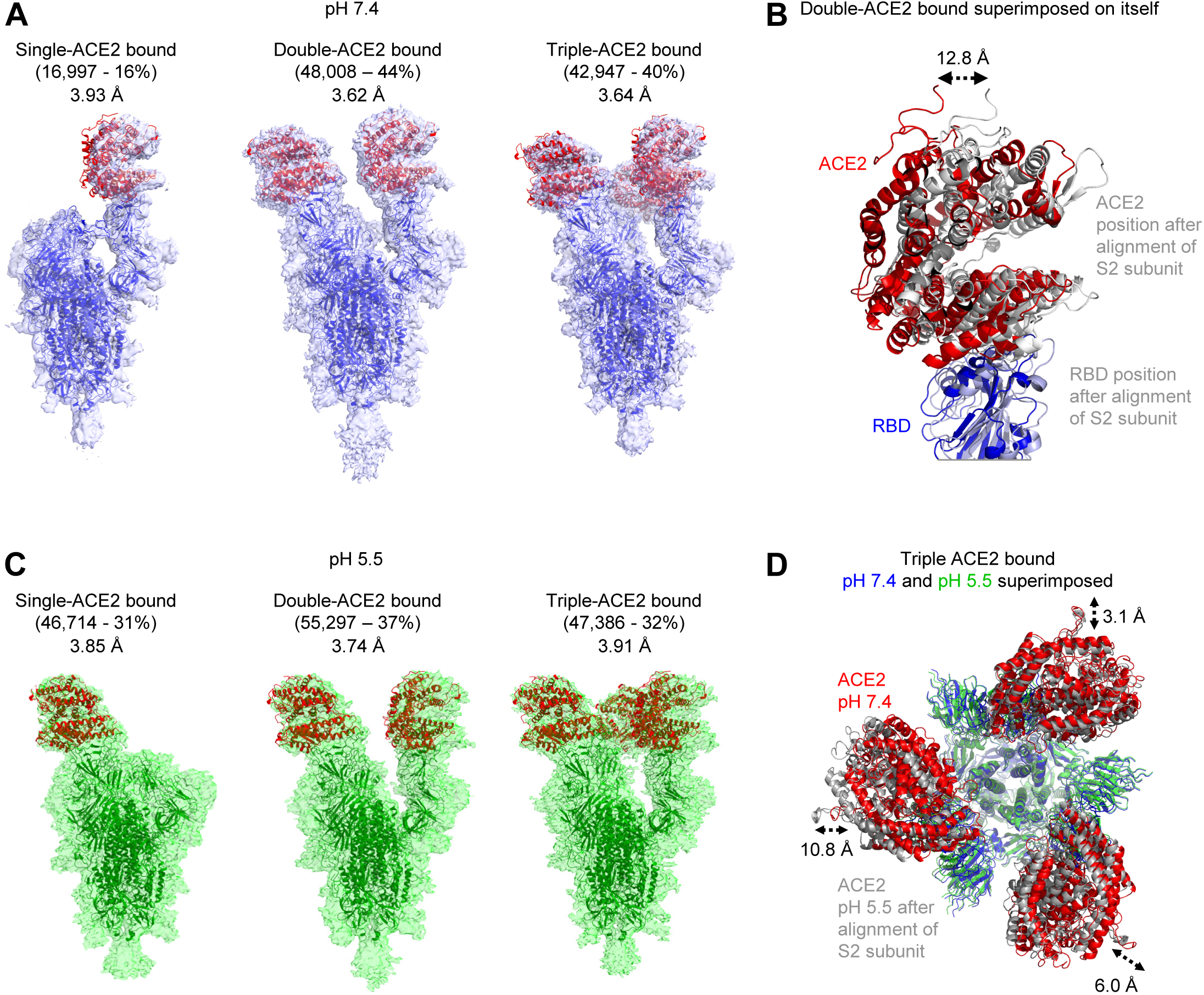
Cryo-EM Structures of SARS-CoV-2 Spike with ACE2 Show Similar Stoichiometries at Serological and Endosomal pH. (**A**) Cryo-EM structures of spike with single-, double-, or triple-bound ACE2 at serological pH. (**B**) Structural comparison of the two ACE2-RBD in the double ACE2-bound structure reveals different tilt angles resulting in as much as a 12.8 displacement as indicated. (**C**) Cryo-EM structure of spike and ACE2 at endosomal pH. (**D**) Comparison of triple CE2-bound kes at serological and endosomal pH. Structures were aligned by S2-subunit superposition and are displayed with the trimer perpendicular to the page and with spike colored according to pH and ACE2 colored red and gray for pH 7.4 and 5.5, respectively.Monomeric ACE2 was used as a ligand in all sample sets. See also Figure S4 and Table S1.

To provide insight into the impact of endosomal pH, we again mixed soluble ACE2 with spike trimer at a 6:1 molar ratio, but this time at pH 5.5, and determined the structure of the complex using cryo-EM. Similar to serological pH, we obtained structures at 3.7-3.9 Å resolution and observed spike to bind ACE2 at stoichiometries of 1:1, 1:2, and 1:3 with prevalences of 31%, 37%, and 32%, respectively (Figure 2C, **Table S1**). We superposed triple-ACE2-bound complexes determined at pH 7.4 and pH 5.5 and observed the membrane-proximal regions of the spike to align closely, while ACE2 molecules showed displacements of 3.1, 6.0, and 10.8 Å (Figure 2D). Overall, structures of the spike with ACE2 showed about equal distribution of single-, double-, and triple-ACE2-bound states at both serological and endosomal pH.

### Ligand-free cryo-EM structures of SARS-CoV-2 spike at low pH

In light of the similarity of ACE2 complexes at pH 7.4 and 5.5 (Figure 2) but substantial differences at these pHs observed for ligand-free spike by negative stain-EM (**Figure S2**), we analyzed the structure of the spike at pH 5.5 by single-particle cryo-EM. We determined a consensus structure from 1,083,554 particles at a resolution of 2.7 Å, in which most of the spike was well resolved, except for a lone RBD for which reconstruction density was poor (Figures 3A **and S5, Table S2**). Analysis of structural heterogeneity in this region (**Videos S1-S4**; **Figure S5C, panel A**) produced six 3D classes ranging in prevalence from 7% to 26% and describing three principal conformations, with the RBD in the up or down position, or without a defined position for this domain (**Figure S5C, panel B**). Interestingly, unlike for ACE2-bound complexes, no double-or triple-RBD-up conformations were observed. Two classes with prevalences of 23% (Conformation 1 – 2.9 Å resolution) and 26% (Conformation 2 – 2.9 Å resolution) corresponded to two different single RBD-up conformations. A third prevalent class representing 10% of the particles had all RBDs down. For all three of these prevalent classes, unlike the consensus structure, density for all RBD domains was well resolved (**Figure S5C, panel C**), indicating multiple different orientations of RBD in the spike at pH 5.5. In the remaining classes, the RBD did not assume a defined position, suggesting RBD mobility at pH 5.5.

To determine how even lower pH affected conformational heterogeneity, and since CR3022 retained binding to spike at pH 5.5 but not at pH 4.5, we sought to obtain a cryo-EM structure of the ligand-free spike at even lower pH. Negative-stain EM analyses indicated a high prevalence of well-formed trimers at both pH 4.5 and 4.0, with some disorder at pH 3.6 (**Figure S2**). We collected cryo-EM datasets at both pH 4.5 and 4.0. Single particle analysis of the pH 4.5 dataset comprising 179,973 particles resolved into an all-RBD-down conformation, and we refined this map to 2.7 Å resolution (Figure 3B **and S5**, **Table S3**); single particle analysis of the pH 4.0 dataset comprising 911,839 particles resolved into a virtually identical all-RBD-down conformation (root-mean square deviation (rmsd) between the two structures of 0.9 Å) (Figure 3C **and S5**). The similarity of the pH 4.5 and pH 4.0 structures indicated spike conformational heterogeneity to be reduced between pH 5.5 and 4.5, and then to remain unchanged as pH was reduced further. The pH 4.0 map was especially well-defined at 2.4 Å resolution (**Table S3**), enabling individual water molecules to be observed (Figure 3D), and we chose the pH 4.0 structure for comparative analysis.

**Figure 3.**
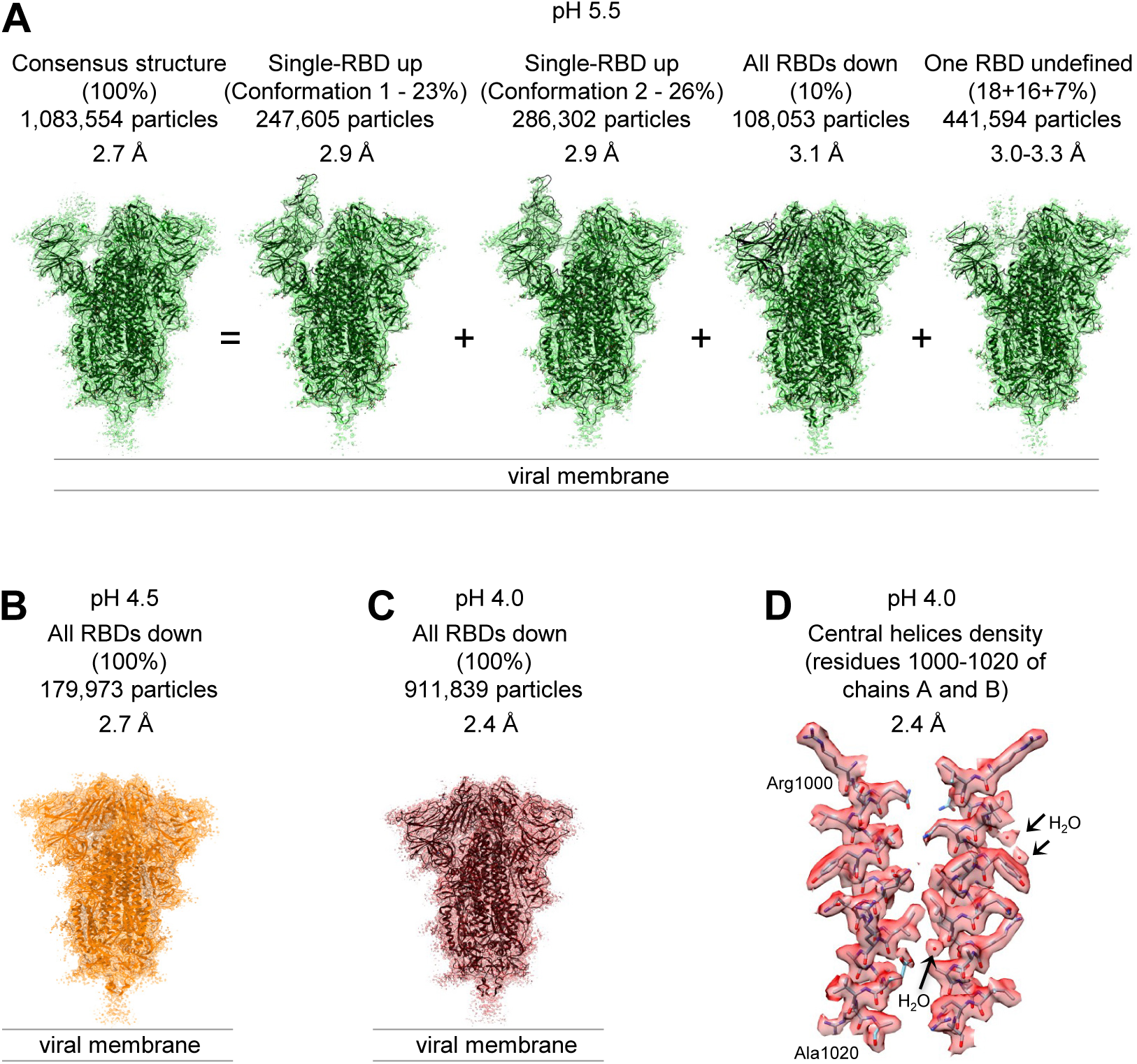
Cryo-EM Analyses Reveal Lower pH to Reduce Spike-Conformational Heterogeneity Culminating in an All RBD-Down Conformation at pH 4.0. (**A**) Structures at pH 5.5 with particle prevalence and resolution of determined structures. (**B**) Structure of spike at pH 4.5. (**C**) Structure of spike at pH 4.5. (**C**) Structure of spike at pH 4.0. (**D**) Example of reconstruction density. A region at the central helices of the pH 4.0 structure is shown with well-defined water molecules. See also Figures S5-S6 and Tables S2 and S3.

### Refolding at spike domain interfaces underlies conformational rearrangement

To identify critical components responsible for the reduction of conformational heterogeneity between pH 5.5 and lower pH and to shed light on the mechanism locking RBDs in the down position, we analyzed rmsds between the pH 5.5 structures and the all-down pH 4.0 conformation with an 11-residue sliding window to identify regions that refold (Figures 4A, **top, and S6**). As each of the protomers in the trimer displayed a different conformation in each of the pH 5.5 structures, we defined protomer B as the one with RBD in the ‘up’ position in each of the single RBD-up conformations, with protomers A-C appearing counter-clockwise when viewed along the trimer 3-fold axis toward the membrane. We observed significant rmsd peaks for short stretches around residue 320 in protomer A only and around residue 525 in protomer B only, and more substantially in a region comprising residues 824-858 (**Figure S6b**). This region, which for reasons described below we named the ‘switch’ region, was fully defined in protomer B and partially resolved in protomer A (residues 824-828 and 848-858) and protomer C (residues 824-841 and 851-858). Notably, this region was almost entirely unresolved in our structures with ACE2 and in most published spike structures (Figure 4A, **bottom**), suggestive of substantial mobility.

**Figure 4.**
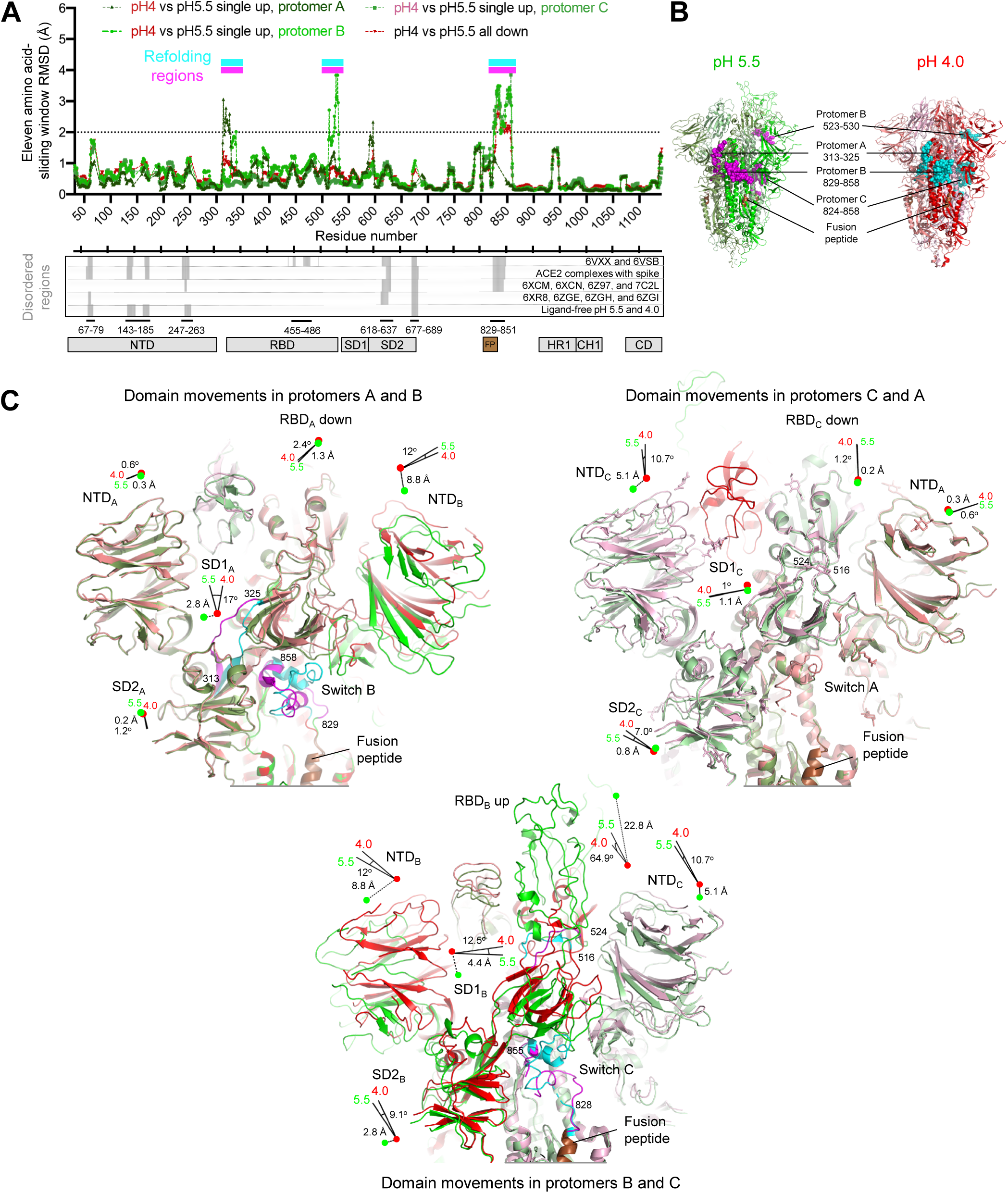
A Switch Domain Mediates RBD Position. (**A**) Identification of refolding regions through rmsd analysis with a 11-residue window (top) and comparison of disordered regions in cryo-EM structures (bottom). (**B**) Refolding regions identified by sliding-window rmsd analysis are highlighted on the pH 5.5 single-up and pH 4.0 structures as spheres and are colored magenta and cyan, respectively. Protomers A, B and C of the pH 5.5 structure are each colored smudge, green or pale green, and the corresponding protomers in the pH 4.0 structure are colored salmon, red or light pink, with fusion peptide colored brown. (**C**) Domain movements between pH 5.5 and 4.0. Three views are shown to depict the movements at the interfaces of protomers A-B, C and C-A. Extent and direction of rotation and displacement are indicated for each domain with vectors and colored dots.Refolding regions are labeled and colored as in (**B**). See also Figures S6-S7 and Table S4.

The asymmetry in distribution of refolding regions in the trimer between single-up and all-down structures (Figure 4B) suggested the ‘up’ RBD to require concerted adjustments throughout the trimer. To delineate these, we determined angles and rigid-body translations between each of the subdomains (**Table S4**) for pH 5.5 single RBD-up and 4.0 all RBD-down structures. For clarity, we specify by subscript the protomer of each subunit or of each residue. Starting with the subdomain 1 of protomer A (SD1_A_) at the entrance loop of protomer A, and moving laterally around the trimer (Figure 4C, **Video S5**), we observed slight refolding in the 313-325_A_ stretch, allowing a 17° rotation of SD1_A_ to accommodate the switch region on the neighboring B protomer (switch B). At pH 5.5, switch B interacted with subdomain 2 of protomer A (SD2_A_) (buried surface area of ~300 Å^2^), and this key inter-protomer contact coupled with SD1_A_ rotation and 2.8-Å translation resulted in the 8.8-Å lateral displacement of N-terminal domain of protomer B (NTD_B_) towards the next RBD-switch (RBD_B_ and switch C). The displaced NTD_B_ induced consecutive shifts of SD2_B_ and SD1_B_ domains, which culminated in the 22.8-Å ‘up’ translation (64.9° rotation) of RBD_B_ versus its down-equivalent.

The ‘up’ positioning of RBD_B_ was accommodated in part by a 5.1 Å mostly downwards displacement of NTD_C_ towards the viral membrane, which – continuing to the next RBD-switch (RBD_C_ and switch A) – induced minor shifts of SD2_C_ and SD1_C_ domains and yielded RBD_C_ and switch A in conformations that closely resembled those of the all-down pH 4.0 structure.

At pH 4.0, each of the RBD-switches closely resembled each other. The most dramatic refolding relative to the switches at pH 5.5 occurred in switch B, where the guanidinium of residue Arg847_B_ swivels over 25 Å from interacting in an inter-protomer manner with SD2_A_ to interacting in an intra-protomer manner with NTD_B_ of the same protomer. This swiveling breaks the coordinated displacements of domains across the protomer-protomer interface, reducing the SD2_A_ interaction with switch B by half (buried surface area of ~160 Å^2^).

Notably, refolding regions were observed to reside at critical inter-protomer contacts or at key joints between domains, especially the SD2 to SD1 joint, which cradles the switch of the neighboring protomer, and the SD1_B_ joint with up-RBD_B_ made up of refolding residues 523-530_B_.

### A pH-dependent switch domain locks spike in down position

The switch domain, which included aspartic acid residues at 830, 839, 843 and 848 and a disulfide linkage between Cys840 and Cys851, was located at the nexus of SD1 and SD2 from one protomer, and HR1 (in the S2 subunit) and NTD from the neighboring protomer. This region showed dramatic conformational changes (Figure 5A). Pairwise rmsd comparisons (Figure 5B) indicated the cryo-EM-determined switch structures to segregate into two conformations: ‘unprotonated-switches’ and ‘protonated-switches’.

**Figure 5.**
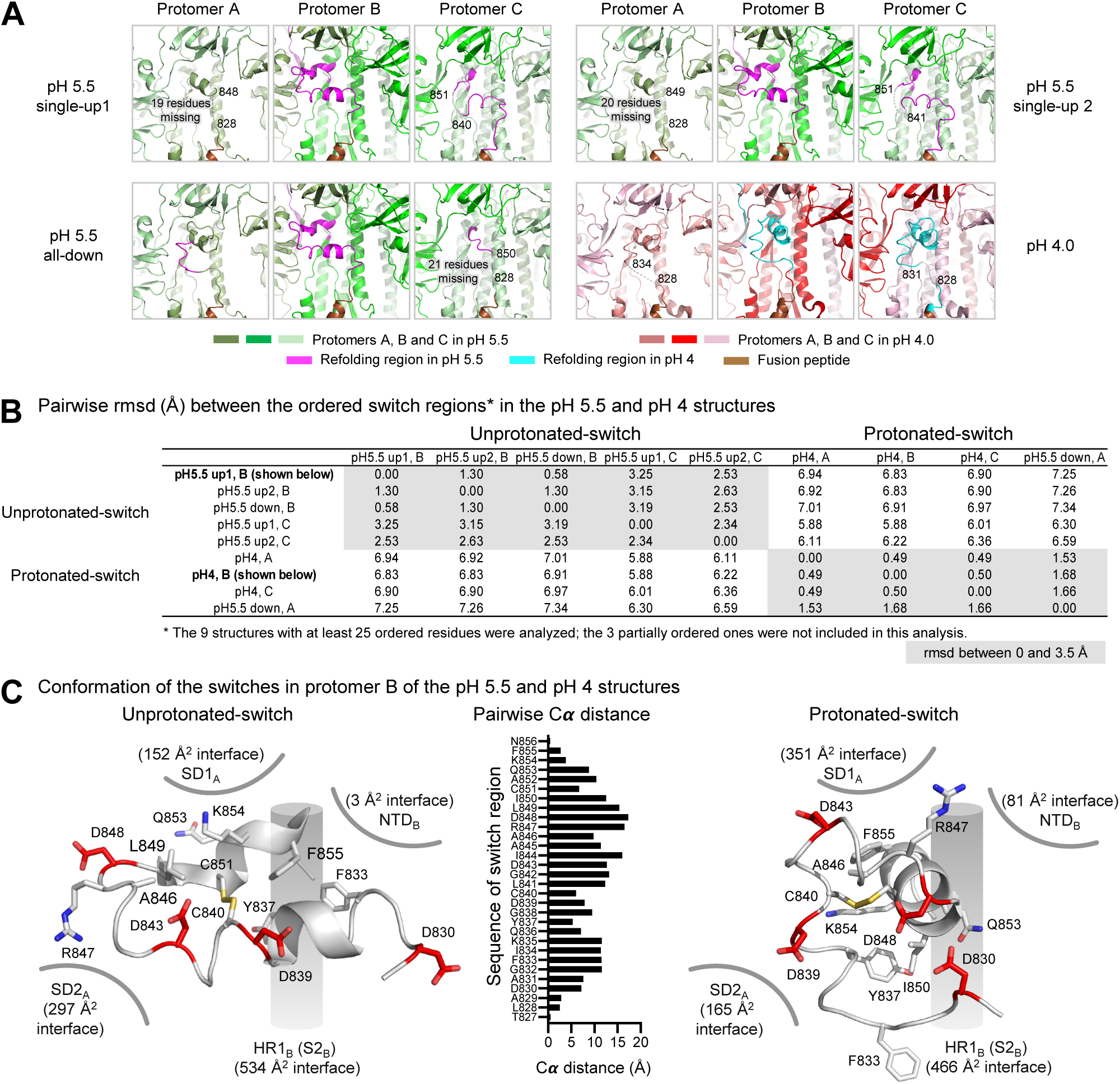
The pH-Switch Domain. (**A**) Switches in the pH 5.5 and pH 4.0 structures. The protomers and switches were colored in Figure 4B. Disordered regions of the switches are shown as gray dashed lines and marked by flanking residue numbers. (**B**) irwise rmsd between switch regions (residues 824-858) from different protomers. Of the 12 protomers determined in this study, ly 9 had at least 25 ordered residues and were included in this pairwise-rmsd analysis; rmsds of less than 3.5 Å shaded grey. itch regions for SARS-CoV-2 spike at higher pH were recently described (Cai et al., 2020; Wrobel et al., 2020) – and these and itch regions from other coronaviruses are analyzed in Figure S7. (**C**) Comparison of the unprotonated and protonated switches. y residues are shown in sticks representation, and Asp and Cys residues are colored red and yellow, respectively. Interactive face areas with surrounding domains indicated. Pairwise Cα-distances between switch residues is shown in the middle. See also Figures S6 and S7, and Tables S5.

Unprotonated-switches were exemplified by switches B and C at pH 5.5 and perhaps best by switch B in the pH 5.5 single-RBD up structure (Figure 5A, C, **left, Video S5**). Continuing from fusion peptide (FP_B_), the N terminus of switch B formed several helical turns (833-842), extending laterally from HR1_B_ to SD2_A_. A turn (843-848) provided extensive contacts with SD2_A_, before returning in helical turns (849-855) back to HR1_B_. Unprotonated-switches were stabilized by a hydrophobic core comprising the disulfide and residues Phe833, Tyr837, Ala846, Leu849, and Phe855 (Figure 5C, **left)**. Notably, all four of the unprotonated-switch aspartic acids faced solvent and appeared to be negatively charged.

Protonated-switches were exemplified by switch A at pH 5.5 and by all switches at pH 4.0 including switch B in the pH 4.0 structure (Figure 5A, C **right, Video S5**). These switches reoriented their N-terminal helical turns to point towards SD1, swiveling the Cα-position of Arg847 over 15 Å to interact with NTD (Figure 5C **and Table S5**) before finishing the rest of the domain with a few helical turns (848-855). Protonated switches were stabilized by a hydrophobic core comprising Tyr837, Ile850, and aliphatic portions of the side chain from Lys854 on one side of the disulfide and Ala846 and Phe855 on the other. Notably, two of the switch domain Asp residues that appeared most likely to be protonated based on hydrogen bonding patterns in the pH 4.0 structure (D830 and D843) also had higher calculated pKas compared to the unprotonated switch conformation, consistent with their observed hydrogen bonds and their apparent protonation at pH 5.5 (Figure 6). Additionally, three Asp residues from the neighboring protomer (D574, D586, and D614) had higher pKas in protonated-switch conformations than in unprotonated-switch conformations. In general, our pKa calculations and structural analyses both indicated increased Asp residue protonation in the protonated-switch conformation, and reflected the expected trend of increasing numbers of protonated Asp or Glu residues at lower pH.

**Figure 6.**
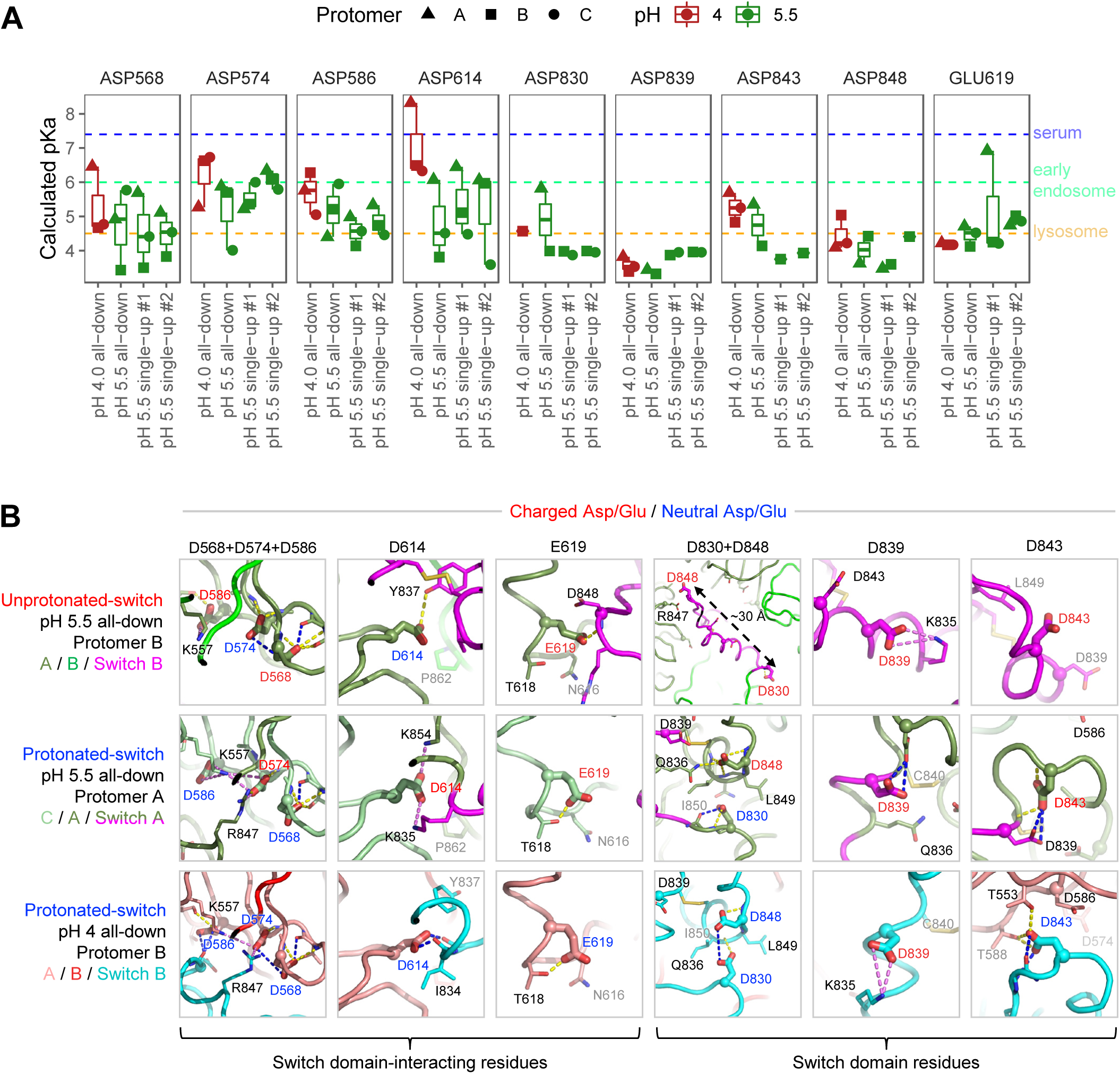
pKa Calculations for the pH-Switch Domain. (**A**) PROPKA-calculated pKas for pH-dependent switch domain idues in the pH 4.0 and 5.5 unliganded spike structures. pKas are plotted for titratable residues within and interacting with the 4-858 pH-dependent switch domain for in each structure, disordered regions excluded. Typical pH values for serum (7.4), early dosome (6.0) and late endosome (4.5) are indicated by dashed lines, each colored as in Figure 1A. (**B**) Close-up views of p/Glu residues in (**A**) from the pH 4.0 and pH 5.5 structures depict changes in chemical environment for each residue between nformations. View angles with respect to superposed structures are the same within each residue column. Switch domain and rounding protomers are colored as indicated at left. Highlighted residues are shown as thick sticks with labels colored based on a-based dominant protonation state at the structure pH: charged Asp/Glu in red, and neutral (protonated) Asp/Glu in blue. sidues within 4 Å are shown as thin sticks. Dashed lines indicate hydrogen bonds (yellow) or salt bridge interactions (violet), th hydrogen bonds requiring carboxylic acid group protonation shown in blue. The pKa shifts between unprotonated-and otonated-switch conformations define a pH-dependent stability gradient that favors the protonated-switch form at lower pHs (Yang & Honig, 1993). However, other factors such as global conformational constraints may also play a role in favoring one conformation over another. See also Figure S7.

Analysis of switch domain conformations and RBD positions (Figure 7A) indicated a concordance between switches interacting with NTD (breaking coordinated interprotomer interactions) and the locking of RBDs in the down position. Thus, at pH 5.5, the unprotonated-switches in protomers B and C interacted with the SD2 domain of the neighboring protomer to transmit lateral displacements of domains. At pH 4.0, the protonated-switches interrupt this interprotomer interaction, resulting in the locking of RBDs in the down position.

**Figure 7.**
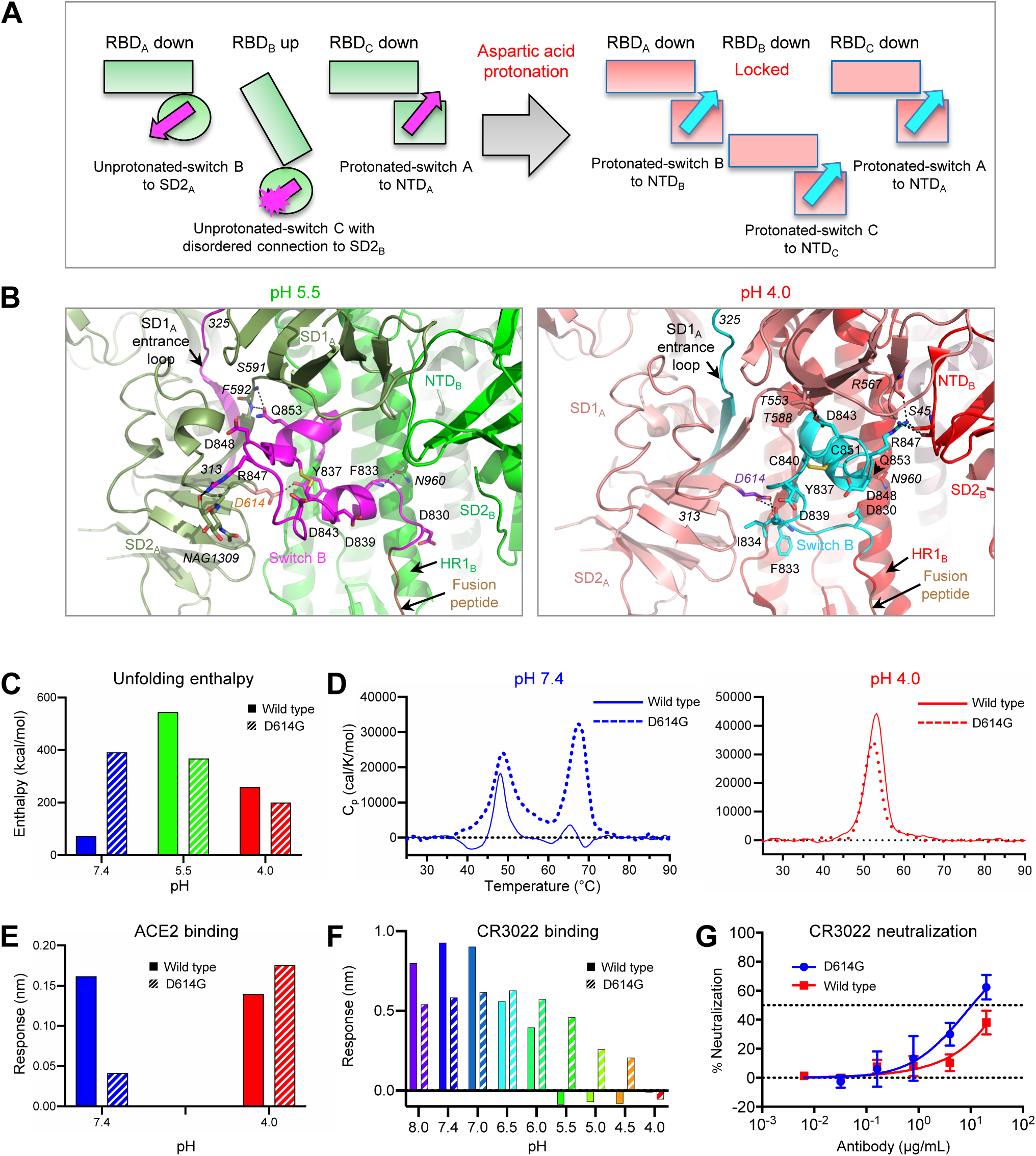
Aspartic Acid Protonation at Low pH Refolds Switch Domain Locking RBDs in the Down Position; an p614Gly Variant Alters SD2-Switch Interactions Leading to Altered ACE2 Interactions and Modestly Impaired Conformational Masking. (**A**) Schematic of the pH-switch locking of RBD in the down position. (**B**) Details of the pH-switch domain. Key residues, including Arg847, Gln853, and Tyr837, switch interactive partners upon refolding. Asp614 is colored orange and purple blue at pH 5.5 and pH 4.0, respectively. Surrounding residues interacting with the switch are labeled in italics, and hydrogen bonds are shown as dashed lines. (**C**) Unfolding enthalpy measured by DSC for the spike and its Asp614Gly variant pH 7.4, 5.5, and 4.0. (**D**) Melting curves at pH 7.4 (left) and 4.0 (right). (**E**) BLI measurements of ACE2 binding to the spike d its Asp614Gly variant at pH 7.4 and 4.0. (**F**) BLI measurements of CR3022 binding to the spike and its D614G variant at different pHs. (**G**) Pseudovirus neutralization of SARS-CoV-2 and its D614G variant by antibody CR3022.

### Impact of Asp614Gly mutation

Analysis of SARS-CoV-2 variant sequences identifies an Asp614Gly mutation to be associated with more transmissible viral variants (Daniloski et al., 2020; Hu et al., 2020; Ke et al., 2020; Korber et al., 2020; Ozono et al., 2020; Yurkovetskiy et al., 2020). Our structures revealed Asp614 to be located at the key interprotomer juncture between SD2 domain and switch, forming a hydrogen bond with Tyr837 of unprotonated-switches (Figure 7B, **left**) and recognizing the backbone carbonyl of Ile 834 in protonated-switches (Figure 7B, **right**). To test the impact of this mutation on unfolding enthalpy, we performed DSC measurements at pH 7.4, 5.5, and 4.0 (Figure 7C). While the enthalpies at pH 5.5 and 4.0 were similar to those of wild-type spike, the unfolding enthalpy at pH 7.4 was dramatically increased, with the appearance of a high melting temperature peak for the variant (Figure 7D), seen for the wild-type only at low pH. The dramatic difference in melting enthalpy demonstrated the substantive energetic effect of altering an interprotomer-switch interface. To test the influence of this increased unfolding enthalpy on ACE2 interaction, we performed biolayer interferometry (BLI) on dimeric ACE2 recognizing spike or Asp614Gly variant. At pH 7.4 we observed higher binding of the wild-type spike than the Asp614Gly variant to dimeric ACE2 (Figure 7E, **left**), consistent with DSC showing more unfolding enthalpy for the variant spike, thereby reducing its ability to bind dimeric ACE2 with avidity.

We next tried to understand the impact of the Asp614Gly mutation on the switch-based mechanism locking RBD in the down position at low pH. As switches with disorder in the region contacting SD2 were associated with “up” RBDs, we hypothesized that mutation of Asp614 to Gly would more closely mimic the loss of interaction between SD2 and switch, as exemplified by RBD_B_ and switch C. Indeed, BLI measurements at pH 4.0 showed dimeric ACE2 to bind the Asp614Gly variant with greater apparent affinity (Figure 7E, **right**), consistent with the higher probability of RBDs adopting the “up” position, and providing an explanation for its increased infectivity (Daniloski et al., 2020; Hu et al., 2020; Korber et al., 2020; Ozono et al., 2020; Yurkovetskiy et al., 2020; Zhang et al., 2020). To test the impact on antibody binding, we used BLI to measure the affinity of CR3022 to spike and Asp614Gly variant. Similar to what we observed with dimeric ACE2, CR3022 bound wild-type spike more tightly than Gly variant at serological pH, with this behavior inverting at low pH where spike folding and switch locking reduced antibody interaction with wild-type but less so with variant spike (Figure 7F). Lastly, we tested the ability of antibody CR3022 to neutralize the Asp614Gly variant, using a pseudovirus format. The Asp614Gly variant showed a modest increase in neutralization sensitivity to CR3022 (Figure 7G), indicating its conformational masking to be mostly intact, with the observed increase in spike binding to CR3022 at low pH perhaps compensating for altered spike interactions with ACE2.

### Conformational masking of SARS-CoV-2 spike

The conformational masking mechanism of immune evasion we delineate here for the SARS-CoV-2 spike involves both serological and endosomal components. At serological pH, the low folding enthalpy of the spike enables antibodies like CR3022 to bind bivalently, with high apparent affinity. Such avidity-based binding would be expected to short-circuit adaptive immune processes of affinity maturation; indeed, analysis of SARS-CoV-2-elicited antibodies indicates RBD-recognizing antibodies to have only a low degree of somatic hypermutation, consistent with impaired maturation (Brouwer et al., 2020; Liu et al., 2020; Robbiani et al., 2020; Rogers et al., 2020; Seydoux et al., 2020).

We demonstrate for CR3022 IgG that its recognition of spike does not impede ACE2 binding, allowing the virus to initiate entry. Once virus binds ACE2, and endosomal entry begins, the pH around the spike would decrease from 7.4 (serum) to ~6.0 (early endosome) and then to 5.5-4.5 (late endosome-early lysosome) (Figure 1). We find this drop in pH has little impact on ACE2 binding (Figure 2), which we observe to maintain nM affinity to spike even at pH 4.5 (**Figure S3B**). We explicitly show >1000-fold relative affinity difference between spike and RBD for CR3022 to occur – not at pH 6.0 (where substantial spike folding occurs) – but at pH 4.5. Thus, it is not the increased unfolding enthalpy of the spike that appears to result in reduced CR3022 affinity, but the pH-switch mediated locking of RBDs in the all-down conformation. We show that between pH 5.5 and 4.5, the spike transitions from a single-RBD-up conformation to a locked all-down state (Figure 3). The all-down state appears to be stably maintained between pH 4.5 and 4.0, and we performed comparative analyses of pH 5.5 and 4.0 structures, revealing the transition to all-down-RBDs to be mediated by a pH-dependent switch, which undergoes dramatic structural refolding (Figures 4-6). We show the switch, located at the nexus of SD1 and SD2 on one protomer and NTD and S2 of another protomer, to be key for controlling the positioning of RBD and in shedding potentially neutralizing antibodies that recognize RBD in the up position.

## Discussion

Viral spikes are prime targets for neutralizing antibody, and many have evolved mechanisms for immune evasion, some of which resemble aspects of the endosomal pH-dependent conformational masking described here. Receptor binding-site masking through endosomal cleavage, for example, occurs with the Ebola virus glycoprotein trimer (Kaletsky et al., 2007), and conformational masking has been previously described for the HIV-1 envelope trimer (Kwong et al., 2002), which is labile and elicits antibodies of little neutralization capacity. With SARS-CoV-2, we delineate conformational masking explicitly here for the non-neutralizing antibody CR3022, which we focused on primarily because of its extensive prior characterization (Huo et al., 2020; Yuan et al., 2020). We anticipate endosomal affinity reduction to apply to all RBD-up recognizing antibodies – including neutralizing antibodies – although this remains to be explicitly shown; we note however that CR3022 has ~100-fold higher affinity to SARS-CoV-1 (Yuan et al., 2020), against which it was originally elicited and which it does neutralize, suggesting its inability to neutralize SARS-CoV-2 stems from a combination of its lower affinity and the endosomal shedding that we describe here.

The functional purpose of the up-down positioning of RBD domains in coronaviruses has been a point of debate, since structures with RBD-up and RBD-down have been determined (Beniac et al., 2006; Gui et al., 2017; Kirchdoerfer et al., 2016; Pallesen et al., 2017; Shang et al., 2020a; Shang et al., 2018; Song et al., 2018; Walls et al., 2016; Yuan et al., 2017). Do the waving RBDs of other coronavirus spikes elicit antibody that is then shed through endosomal entry mechanisms along the lines that we outline for SARS-CoV-2? We note that the switch domains from bat RaTG13 and SARS-CoV-1 are virtually identical in sequence to that of SARS-CoV-2, and aspartic acids residues are mostly conserved in MERS (**Figures S7**), potentially indicating the switch-based all-RBD-down locking strategy of immune evasion described here to enable other coronaviruses that utilize endosomal entry to avoid neutralization by RBD-up-recognizing antibody.

The critical switch region (residues 824-858) displays remarkable structural diversity within coronaviruses, segregating into three structural clusters (**Figure S7**). Each of the structures within these clusters generally comprises two helices, linked by a disulfide, in distinct orientations relative to each other and to the surrounding domains. The structural diversity of the switch region, defined here for SARS-CoV-2 along its endosomal entry pathway and recently at higher pH (Cai et al., 2020; Wrobel et al., 2020), provides a further example of how type 1 fusion machines can use structural rearrangement not only to merge membranes (e.g. transitioning from prefusion to intermediate to postfusion states) but to evade potential neutralizing antibodies that recognize the prefusion state.

## Acknowledgements

We thank S. Goff for discussions on viral variants and entry mechanism, R. Grassucci, Y.-C. Chi and Z. Zhang from the Cryo-EM Center at Columbia University for assistance with cryo-EM data collection, M.G. Joyce for antibody CR3022, J.S. McLellan for spike expression vector, E.H. Zhou for assistance with movies, and members of the Virology Laboratory and Vector Core, Vaccine Research Center, for discussions and comments on the manuscript. Support for this work was provided by the Intramural Research Program of the Vaccine Research Center, National Institute of Allergy and Infectious Diseases (NIAID), Federal funds from the Frederick National Laboratory for Cancer Research under Contract HHSN261200800001E (A.S., T.S., Y.T.). Cryo-EM data for the spike-ACE2 complexes were collected at Columbia University Cryo-EM Center at the Zuckerman Institute, and at the National Center for CryoEM Access and Training (NCCAT) and the Simons Electron Microscopy Center located at the New York Structural Biology Center, supported by the NIH Common Fund Transformative High Resolution Cryo-Electron Microscopy program (U24 GM129539,) and by grants from the Simons Foundation (SF349247) and NY State Assembly. Cryo-EM datasets for individual spike proteins were collected at the National CryoEM Facility (NCEF) of the National Cancer Institute. This research was, in part, supported by the National Cancer Institute’s National Cryo-EM Facility at the Frederick National Laboratory for Cancer Research under contract HSSN261200800001E. We are especially grateful to U. Baxa, A. Wier, M. Hutchison, and T. Edwards of NCEF for collecting cryo-EM data and for technical assistance with cryo-EM data processing. Frederick Research Computing Environment (FRCE) high-performance computing cluster was used for processing cryo-EM datasets of individual spike proteins.

## Author Contributions

Y.T. and T.Z. determined ligand-free spike structures at pH 5.5, 4.5, and 4.0; A.S.O. produced spike and Asp614Gly variant and performed DSC; J.G. M.R. and G.C. determined spike-ACE2 structures; G.-Y.C. carried out informatics analyses; P.S.K. performed SPR; A.N. carried out BLI; J.M.S. calculated pKa; A.S. performed ITC; P.W. preformed neutralization assessments; J.B., W.S., I.T.T., B.Z. provided reagents; J.C.B. analyzed switch mechanics; T.S. prepared ligand-free cryo-EM specimens; M.S. produced spike expression vectors; J.S. assisted with entry mechanism; S.W. assisted with manuscript preparation; R.A.F. supervised pKa calculations; D.D.H. supervised neutralization; J.R.M. supervised reagents and analyses, L.S. supervised SPR and cryo-EM studies with ACE2; P.D.K. oversaw the project and –with T.Z., Y.T., A.S.O., J.G., P.S.K. A.N., A.S., P.W., W.S., B.Z., G.-Y.C., J.M.S., S.W., and L.S. – wrote the manuscript, with all authors providing revisions and comments.

## Competing interest declaration

The authors declare no competing interest.

## STAR⍰METHODS

### RESOURCE AVAILABILITY

#### Lead Contact

Further information and requests for resources and reagents should be directed to and will be fulfilled by the Lead Contact, Peter D. Kwong.

#### Materials Availability

This study did not generate new unique reagents.

#### Data and Code Availability

Cryo-EM structure coordinates and electron density maps for the SARS-CoV-2 spike ligand free and ACE2 complexes are in the process of being deposited with the Protein Data Bank and Electron Microscopy Data Bank.

### EXPERIMENTAL MODEL AND SUBJECT DETAILS

#### Cell Lines

FreeStyle 293-F (cat# R79007) and Expi293F cells (cat# A14528; RRID: CVCL_D615) were purchased from ThermoFisher Scientific Inc. FreeStyle 293-F cells were maintained in FreeStyle 293 Expression Medium, while Expi293F cells were maintained in Expi Expression Medium. The above cell lines were used directly from the commercial sources and cultured according to manufacturer suggestions. HEK293T (cat# CRL-11268) and Vera E6 cells (cat# CRL-1586) were purchased from ATCC and were maintained and used according to manufacturer instructions.

### METHOD DETAILS

#### Production of spike, ACE2 receptor and antibodies

SARS-CoV-2 spike (Wrapp et al., 2020) and its D614G mutant were expressed by transient transfection in 293 Freestyle cells. Briefly, 1 mg of DNA was transfected into 1L of cells using Turbo293 transfection reagent, and the cells were allowed to grow at 37°C for 6 days. Following expression, the supernatant was cleared by centrifugation and filtration, and then incubated with cOmplete His-Tag Purification resin. The resin was washed with PBS containing increasing concentrations of imidazole, and the protein eluted in 20 mM Tris pH8.0, 200 mM NaCl, 300 mM Imidazole. HRV3C protease was added at a 1:20 mass ratio and incubated overnight at 4 °C to cleave the purification tags. The protein was then applied to a Superdex 200 column in PBS, after which the spike containing fractions were pooled and concentrated to 1 mg/ml. Single chain Fc tagged RBD and NTD domains were expressed in the same manner, and purified using capture by Protein A resin, followed by cleavage of the tag using HRV3C (Zhou et al., 2020b) and gel filtration.

Human ACE2 proteins were prepared in monomeric form (residues 1-620) and in dimeric form (residues 1-740). The expression plasmids were constructed and the protein purified as described previously (Zhou et al., 2020b). Briefly, DNA sequence encoding monomeric or dimeric ACE2 was synthesized and cloned into a plasmid with an HRV3C cleavage site, monomeric Fc tag and 8xHisTag at the 3’-end. The proteins were expressed by transient transfection of 293F cells and purified from a Protein A column. The tag was removed by overnight HRV3C digestion at 4 °C. The proteins were further purified with a Superdex 200 16/60 column in 5 mM HEPES, pH7.5 and 150 mM NaCl.

For antibody preparation, DNA sequences of antibody CR3022 (Yuan et al., 2020) heavy and light chains were cloned into the pVRC8400 vector, as described previously (Wu et al., 2011), expressed and purified as described (Zhou et al., 2020b). The Fab fragments were generated by overnight digestion with Endoproteinase LysC (New England Biolabs) at 37 °C and purified by protein A column to remove uncut IgG and Fc fragments.

#### Differential scanning calorimetry

DSC analyses were performed using a Microcal VP-Capillary DSC. The proteins were diluted to 0.25 mg/ml in 1X PBS or various solutions containing a final concentration of 100 mM buffer and 200mM NaCl. The buffers used were: pH 4.0-pH 5.5, Sodium Acetate; pH 6.0-pH 6.5, MES; pH 7.0, HEPES; pH 7.4, 1X PBS; pH 8.0-pH 8.5, Tris; and pH 9.0, Sodium Borate. The proteins were scanned at 1 °C per minute from 25 – 90 °C using a filter period of 25 s. Data were analyzed using the Origin based Microcal DSC Automated Analysis software, where baselines were subtracted, and peak area and T_m_ calculated. The melting curves and unfolding enthalpy graphs were made in Excel.

#### Isothermal titration calorimetry

Calorimetric titration experiments were performed at 25 °C using a VP-ITC microcalorimeter from MicroCal⁄Malvern Instruments (Northampton, MA, USA). The spike protein, ACE2 and Fab of CR3022 were prepared and exhaustively dialyzed against PBS, pH 7.4, prior to the experiments. Any dilution steps prior to the experiments were made using the dialysate to avoid any unnecessary heats of dilution associated with the injections. All reagents were thoroughly degassed prior to the experiments. For the direct determination of the binding to the spike protein, the solution containing either ACE2 or CR3022(Fab) was added stepwise in 10 µL aliquots to the stirred calorimetric cell (~ 1.4 ml) containing spike protein at 0.4 – 0.5 µM (expressed per trimer). The concentration of titrant in the syringe was 12 – 14 µM for both ACE2 and CR3022(Fab). The effect of CR3022 on ACE2 binding to spike protein was studied by first titrating the spike protein with CR3022 until complete saturation was reached, and then performing a complete titration of the complex with ACE2. Despite the thorough dialysis, the heat of dilution/injection associated with the injection of ACE2 into the complex was considerable during the course of the titration and needed to be accounted for in the analysis. The heat evolved upon each injection was obtained from the integral of the calorimetric signal and the heat associated with binding was obtained after subtraction of the heat of dilution. The enthalpy change, *ΔH*, the association constant, *K_a_*, and the stoichiometry, *N*, were obtained by nonlinear regression of the data to a single-site binding model using Origin with a fitting function made inhouse. Gibbs energy, *ΔG*, was calculated from the binding affinity using *ΔG* = -*RT*ln*K_a_*, (*R* = 1.987 cal/(K × mol)) and *T* is the absolute temperature in kelvin). The entropy contribution to Gibbs energy, *-TΔS*, was calculated from the relation *ΔG* = *ΔH* -*TΔS*.

#### SPR binding experiments

SPR binding experiments were performed using a Biacore T200 biosensor, equipped with a Series S SA chip. The running buffer varied depending on the pH of the binding reaction; experiments at pH 7.4 were performed in a running buffer of 10 mM HEPES pH 7.4, 150 mM NaCl, 0.2 mg/ml BSA and 0.01% (v/v) Tween-20; at pH 5.5 experiments were performed in 10 mM sodium acetate pH 5.5, 150 mM NaCl, 0.2 mg/ml BSA and 0.01% (v/v) Tween-20; and at pH 4.5 in 10 mM sodium acetate pH 4.5, 150 mM NaCl, 0.2 mg/mL BSA and 0.01% (v/v) Tween-20. All measurements were performed at 25 °C.

Biotinylated spike and RBD were captured over independent flow cells at 700-1000 RU and 150 RU respectively for both the CR3022 IgG and the dimeric ACE2 binding experiments. To avoid the difficulty in surface regeneration that arises with slow dissociation, we used single-cycle kinetics binding experiments. CR3022 IgG was tested at analyte concentrations 36-1.33 nM prepared in running buffer at each pH, using a three-fold dilution series. In addition, CR3022 IgG was tested over the spike at higher analyte concentrations ranging 108-4 nM, 360-13.33 nM and 1000-37.04 nM at pH 4.5, only to confirm the absence of binding to the spike at pH 4.5. Dimeric ACE2 was tested at 90-3.33 nM prepared in running buffer at each pH, using a three-fold dilution series. Binding over the spike or RBD surface as well as over a streptavidin reference surface was monitored for 120 s, followed by a dissociation phase of 120-900 s depending on the interaction at 50 μl/min. Four blank buffer single cycles were performed by injecting running buffer instead of Fab to remove systematic noise from the binding signal. The data was processed and fit to 1:1 single cycle model using Scrubber 2.0 (BioLogic Software).

#### Cryo-EM structures of ACE2-spike complexes

SARS-CoV-2 spike was incubated with 3-fold molar excess of ACE2 receptor with a final trimer concentration of 1 mg/ml in either PBS, pH 7.4, or 10 mM sodium acetate, pH 5.5, with 150 mM NaCl. The samples (2 µl) were vitrified using a Leica EM GP and Vitrobot Mark IV plunge freezers on glow-discharged carbon-coated copper grid (protochip, CF 1.2/1.3). Data were collected on a 300 kV Titan Krios equipped with a Gatan K3-BioQuantum direct detection device using Leginon software (Suloway et al., 2005). The total dose was fractionated for 2 s over 40 raw frames. Motion correction, contrast transfer function (CTF) estimation, particle picking with topaz (Bepler et al., 2019) and extraction, 2D classification, ab initio model generation, 3D refinements and local resolution estimation were carried out in cryoSPARC 2.14 (Punjani et al., 2017). We note that some classes of unbound spike were also observed in both datasets however particle picking was optimized for complexes so the fraction was low. The 3D reconstructions were performed using C1 symmetry for all complexes as the ACE2-RBD region showed flexibility that prohibited typical symmetry operations in the triple-bound complexes. However, the RBD-ACE2 region was assessed in greater detail through focused refinement following particle expansion with C3 symmetry applied to the pH 7.4 triple bound reconstruction. This RBD-ACE2 model was then used as a reference structure for refinement of all other ACE2-bound models.

The coordinates of SARS CoV-2 spike ectodomain structures, PDB entries 6VXX and 6M0J (Walls et al., 2020), were employed as initial models for fitting the cryo-EM map of theACE2 bound structures. Manual and automated model building were iteratively performed using Coot (Emsley and Cowtan, 2004) and real space refinement in Phenix to accurately fit the coordinates to the electron density map. Molprobity (Davis et al., 2004) was used to validate geometry and check structure quality. UCSF ChimeraX (Goddard et al., 2018) was used for map-fitting cross correlation calculation (Fit-in-Map tool) and for figure preparation.

#### Negative-stain electron microscopy

The following buffers were used to study SARS-CoV-2 S conformation at different pH: 0.1 M sodium acetate (pH 3.6–5.5), PBS (pH 7.4), 0.1 M Trizma-HCl (pH 8.8). A sample of SARS-CoV-2 S with a concentration of 1 mg/ml was diluted with the target buffer 10 times, and the diluted sample was incubated on ice for 15 min. Immediately before negative staining, the sample was further diluted 5 times with the following buffer: 10 mM sodium acetate, 150 mM NaCl (for pH 3.6–5.5); 10 mM HEPES, 150 mM NaCl (for pH 7.4); 10 mM Trizma-HCl, 150 mM NaCl (for pH 8.8). A 4.7-µl drop of the diluted sample was applied to a glow-discharged carbon-coated copper grid for 10-15 s. The drop was removed with filter paper, and the grid was washed by applying consecutively three 4.7-µl drops of the buffer used for diluting the sample and removing them with filter paper. Protein molecules adsorbed to the carbon were negatively stained by applying consecutively three 4.7-µl drops of 0.75% uranyl formate in the same manner. The grid was air-dried and screened for staining quality and particle density using a Hitachi H-7650 transmission electron microscope (TEM). Datasets were collected using an FEI T20 TEM equipped with an Eagle CCD camera. The microscope was operated at 200 kV, the pixel size was 2.2 Å (nominal magnification: 100,000), and the defocus was −1.2 µm. SerialEM (Mastronarde, 2005) was used for data collection. Particles were picked automatically and extracted into 160×160-or 192×192-pixel boxes using in-house written software (YT, unpublished). 2D classification was performed using Relion 1.4 and Relion 3.0 (Scheres, 2012).

#### Cryo-EM specimen preparation and data collection of individual spikes

A sample of SARS-CoV-2 S in PBS with a protein concentration of 1 mg/ml was diluted to 0.5 mg/ml using 0.2 M sodium acetate, pH 4.0 or pH 5.5 (final sodium acetate concentration: 0.1 M). Separate measurements with a pH meter confirmed that combining equal volumes of PBS and 0.2 M sodium acetate, pH 4.0 or pH 5.5, produces solutions with pH 4.0 and pH 5.5, respectively. Quantifoil R 2/2 gold grids were used for specimen preparation. The grids were glow-discharged using a PELCO easiGlow device (air pressure: 0.39 mBar, current: 20 mA, duration: 30 s) immediately before vitrification. Cryo-EM grids were prepared by plunge-freezing in liquid ethane using an FEI Vitrobot Mark IV plunger with the following settings: chamber temperature of 4°C, chamber humidity of 95%, blotting force of −5, blotting time of 5 s, and drop volume of 2.7 µl. Datasets were collected at the National CryoEM Facility (NCEF), National Cancer Institute, on a Thermo Scientific Titan Krios G3 electron microscope equipped with a Gatan Quantum GIF energy filter (slit width: 20 eV) and a Gatan K3 direct electron detector. Four movies per hole were recorded in the counting mode using Latitude software. The dose rate was 13.4 e^-^/s/pixel.

#### Cryo-EM data processing and structural refinement for individual spikes

Each dataset was divided into subsets which were initially processed independently in parallel using Frederick Research Computing Environment (FRCE) computing cluster and later combined for the final refinement. Movie frame alignment was performed using MotionCorr2 (Zheng et al., 2017). Ctffind4 was used to determine the parameters of CTF (Rohou and Grigorieff, 2015). The remaining processing steps were performed using Relion 3.0 (Scheres, 2012) unless otherwise stated. For spike at pH 4.0, a small particle set was selected manually and used to obtain 2D classes which were utilized as templates to select a larger set of particles. An initial 3D model was obtained using EMAN 2.1 (Tang et al., 2007) from the 2D classes generated from this extended particle set. This 3D model was then subjected to 3D auto-refinement, and the resulting map was used to generate low-pass filtered picking templates for the entire dataset. For spike at pH 5.5, particle picking was performed with cryOLO 1.5 (Wagner et al., 2019) using a general network model, and an initial 3D model was obtained with EMAN 2.1 from a subset of resulting 2D classes. The following steps included rounds of 3D classification, 3D auto-refinement, CTF refinement, and particle polishing. Map resolutions were calculated using the gold-standard approach (Henderson et al., 2012) at the FSC curve threshold of 0.143. ResMap 1.1.4 was used to asses local resolution (Kucukelbir et al., 2014). Local map sharpening was performed using phenix.auto_sharpen (Terwilliger et al., 2018). SPIDER 22.1 was used for map conversion and resizing (Frank et al., 1996). Correlations between cryo-EM maps and atomic models were assessed using phenix.mtriage (Afonine et al., 2018). UCSF Chimera was used for docking and visualization (Pettersen et al., 2004). Despite the fact that C3 symmetry was imposed during the reconstruction of spike for the pH 4.0 dataset, the resulting map displayed some asymmetrical features in some regions, such as that around residue 830. Therefore, the three chains of the atomic model were built and refined individually. The coordinates of SARS CoV-2 spike ectodomain structures, PDB entries 6VXX and 6VYB, were used as initial models for fitting the cryo-EM map of the spike structures at pH 4.0 and pH 5.5 structures. Iterative model building and real space refinement were carried out using Coot (Emsley and Cowtan, 2004) and Phenix to accurately fit the coordinates to the electron density map. Molprobity (Davis et al., 2004) was used to validate geometry and check structure quality.

#### 3D variability analysis of cryo-EM structures of individual spikes and analysis of conformations of the RBD

For 3D variability analysis, a subset of 100,000 particles randomly selected from the final particle set at pH 5.5 was exported into cryoSPARC 2.15 (Punjani et al., 2017), and a homogeneous refinement was performed without imposing symmetry. The 3D variability analysis was set up to use three eigenvectors of the 3D covariance, and 20 frames were used for visualization of results. The eigenvectors describing movements of the RBD were identified via examining the resulting volume series and corresponding variability movies (Videos S1-4).

The structural heterogeneity of the consensus pH 5.5 map in the RBD region was analyzed using local 3D classification. To obtain an accurate mask encompassing the conformational space of the dynamic RBD, the four 3D variability volumes corresponding to the beginning and the end of the trajectories defined by eigenvectors 0 and 2 were first aligned to the consensus cryo-EM map. For each of the four volumes, the density corresponding to the dynamic RBD was isolated by performing volume segmentation in UCSF Chimera (Pettersen et al., 2004). These RBD sub-volumes were added together, and a soft mask was created from the resulting composite volume by low-pass filtering the density to 15 Å, extending the resulting volume by 2 pixels, and adding a soft edge of 5 pixels using relion_mask_create. Local 3D classification of the consensus dataset within this mask was performed without particle alignment in Relion 3 (Scheres, 2012), followed by global 3D refinement of each of the resulting six maps.

#### Identification of SARS-CoV-2 spike refolding regions between pH 5.5 and pH 4.0 structures

We used a sliding window of 11 amino acids and 21 amino acids respectively to align and calculate backbone (C, Ca, O, N) rmsd values between the pH 4 structure (protomer B) and pH 5.5 single-RBD-up or pH 5.5 all-RBD-down structures, respectively, using PyMol (Version 2.3.4). Calculation was omitted if the specified residue range had less than 22 backbone atoms. The average rmsd values of pH 5.5 single-RBD-up conformation 1 and conformation 2 were reported for pH 5.5 single-RBD-up analysis. The refolding regions were defined as residues with greater than 2-Å rmsd. Refolding regions with more than one consecutive residue were further considered, and single residue gaps were ignored when determining the residue ranges. Manual inspection revealed 11 amino acid-window to correspond better with domain movements. Therefore, the results from only 11 amino acid-window analysis were reported.

#### Clustering of coronavirus spike structures based on the switch region

The coronavirus spike structures were obtained from PDB using sequence similarity search against SARS-CoV-2 spike protein with the default parameters. After manual examinations, structures that were not coronavirus spike trimers were excluded. For the rest of the structures, the sequences were aligned using ClustalW (Larkin et al., 2007) and chains with at least 70% of the residues determined of the switch region (residues 824-858, SARS-CoV-2 numbering) were further considered. The structures were clustered using the hclust function implemented in statistical package R based on the pairwise backbone rmsd distances calculated with the rms_cur function in PyMOL after the switch regions were aligned.

#### Bio-layer interferometry (BLI)

A FortéBio Octet HTX instrument (FortéBio) was used to assess binding over a wide pH range. Experiments were setup in tilted black 384-well plates (FortéBio) in 10mM of the corresponding buffer, plus 150mM NaCl, 0.02% Tween20, 0.1% BSA and 0.05% sodium azide. Buffers used for pH 8.0 to 4.0 are as described above in the DSC section. Plates were agitated at 1,000 rpm, and the temperature was set to 30°C. Anti-human IgG Fc capture biosensors (FortéBio) were used to immobilize 300nM CR3022 IgG or dimeric ACE2-Fc for 150 seconds at pH 7.4. Following loading of CR3022 IgG, sensors were placed in the pH 7.4 buffer for 30 seconds and then equilibrated in the respective pH buffer for 180 seconds. Binding was measured for 180 seconds in 200 nM spike or D614G mutant. Binding quantification (Figure 7d, e) was performed using the response value (nm) in the last second of the association step. Dissociation in the respective buffer was recorded for 300 seconds.

#### pKa calculations

Individual residue pKas were calculated for the pH 4.0 all-down, pH 5.5 all-down, and pH 5.5 single-up (conformations 1 and 2) structures using PROPKA (Olsson et al., 2011; Sondergaard et al., 2011). For residues in the chain B 830-855 switch domains and titratable residues within 5Å of the switch domain, pKa data were analyzed and plotted using R (https://www.R-project.org/) in RStudio (http://www.rstudio.com/) with the ggplot2 library (Wickham, 2016) and structural figures were made using PyMOL.

#### Pseudovirus construction and neutralization assessment

Recombinant Indiana vesiculovirus (rVSV) expressing SARS-CoV-2 spike was generated as previously described (Nie et al., 2020; Whitt, 2010). HEK293T cells were grown to 80% confluency before transfection with pCMV3-SARS-CoV-2-spike (kindly provided by Peihui Wang, Shandong University, China) or the D614G variant (constructed by site-directed mutagenesis) using FuGENE 6 (Promega). The next day, medium was removed and VSV-G pseudotyped ΔG-luciferase (G*ΔG-luciferase, Kerafast) was used to infect the cells in DMEM at an MOI of 3 for 1 h before washing the cells with 1X DPBS three times. DMEM supplemented with 2% fetal bovine serum and 100 I.U./mL of penicillin and 100 μg/mL of streptomycin was added to the inoculated cells. The supernatant was harvested the following day and clarified by centrifugation at 3000 rpm for 10 min before aliquoting and storing at −80°C.

Neutralization assays were performed by incubating pseudoviruses with serial dilutions of antibodies and scored by the reduction in luciferase gene expression (Liu et al., 2020). In brief, Vero E6 cells (ATCC) were seeded in a 96-well plate at a concentration of 2 × 10^4^ cells per well. Pseudoviruses were incubated the next day with serial dilutions of the antibodies in triplicate for 30 min at 37°C. The mixture was added to cultured cells and incubated for an additional 24 h. The luminescence was measured by Britelite plus Reporter Gene Assay System (PerkinElmer). IC_50_ was defined as the dilution at which the relative light units (RLUs) were reduced by 50% compared with the virus control wells (virus + cells) after subtraction of the background RLUs in the control groups with cells only. The IC_50_ values were calculated using non-linear regression in GraphPad Prism 8.

### QUANTIFICATION AND STATISTICAL ANALYSIS

The BLI and DSC data were analyzed and plotted using Excel and GraphPad Prism. The SPR data were processed and fit using Scrubber 2.0 (BioLogic Software). Cryo-EM data were processed and analyzed using CryoSparc and Relion. Cryo-EM structural statistics were analyzed with Phenix and Molprobity. Statistical details of experiments are described in Method Details or figure legends.

## Supplemental Video Legends

Video S1. A side-view movie illustrating the trajectory of the 3D covariance described by eigenvector 0 in 3D variability analysis of individual spike at pH 5.5 (see Methods for a detailed description). A ratcheting motion of one NTD domain results in increased mobility of the corresponding RBD. A corresponding top view is presented in Video S2.

Video S2. A top-view movie illustrating the trajectory of the 3D covariance described by eigenvector 0 in 3D variability analysis of individual spike at pH 5.5 (see Methods for a detailed description). The RBD is up and alternates between two positions. A corresponding side view is presented in Video S1.

Video S3. A side-view movie illustrating the trajectory of the 3D covariance described by eigenvector 2 in 3D variability analysis of individual spike at pH 5.5 (see Methods for a detailed description). A ratcheting motion of one NTD domain results in increased mobility of the corresponding RBD. A corresponding top view is presented in Video S4.

Video S4. A top-view movie illustrating the trajectory of the 3D covariance described by eigenvector 2 in 3D variability analysis of individual spike at pH 5.5 (see Methods for a detailed description). The RBD alternates between up and down positions. A corresponding side view is presented in Video S3.

Video S5. pH-dependent domain movements in the SARS-CoV-2 spike and pH-switch refolding.

## Notes

### Competing Interest Statement

The authors have declared no competing interest.

