## Supplemental Material for "Cryo-EM Structures Delineate a pH-Dependent Switch that Mediates Endosomal Positioning of SARS-CoV-2 Spike Receptor-Binding Domains"

**A**

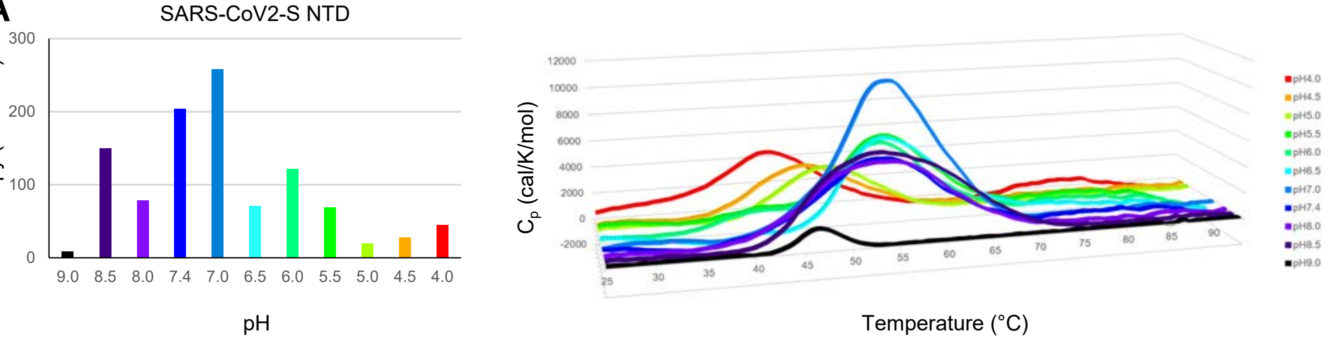


**B**

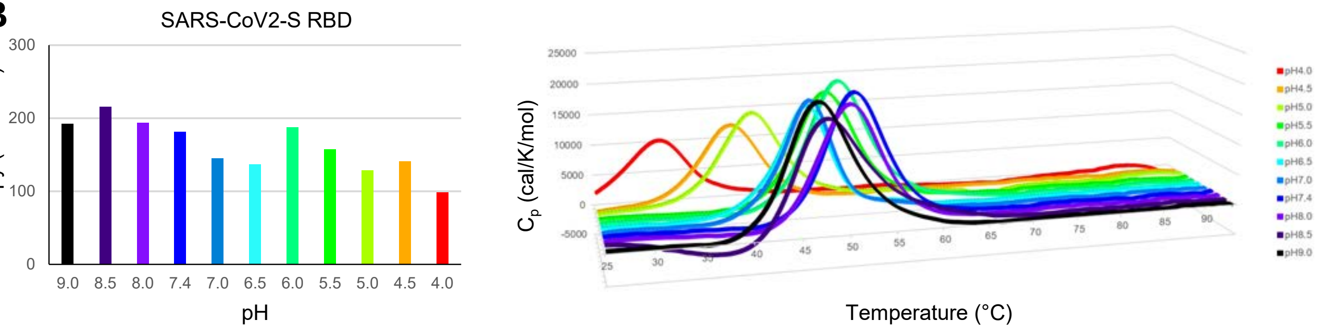


**C**

| | $T_m$ (°C) | Molar unfolding enthalpy (cal/mol) | Mass unfolding enthalpy (cal/g) | Average protein unfolding enthalpy* (cal/g) | Percent of normalized protein (%) |
|---|---|---|---|---|---|
| Spike | 48 | 73,330 | 0.531 | 5.26 | ~10 |
| NTD | 55 | 204,085 | 5.831 | 6.10 | >90 |
| RBD | 52 | 181,051 | 5.173 | 5.74 | >90 |

* Calculated by $\Delta H = \Delta H_{ref} + \Delta C_p(T-T_{ref})$ where $T_{ref}$=60°C, $\Delta H_{ref}$=6.7cal/g, and $\Delta C_p$=0.12cal/K/g

**Figure S1. Unfolding Enthalpy of SARS-CoV-2 NTD and RBD as a Function of pH, Related to Figure 1.** (**A**) DSC denaturation curves for SARS-CoV-2 NTD. (**B**) DSC denaturation curves for SARS-CoV-2 RBD. (**C**) Calculations of normalized unfolding enthalpy relative to that of an average globular protein. The enthalpy of unfolding of the spike protein is considerably smaller than expected at pH 7.4 and above. The thermal unfolding of the spike at pH 7.4 has its main transition centered at 48 °C with an overall enthalpy of unfolding of 73.3 kcal/mol or 0.531 cal/g, which is about an order of magnitude smaller than expected for a protein of this size. In comparison, the expected unfolding enthalpy for the average globular protein is about 5.3 cal/g at this temperature (obtained after extrapolation from 60 °C, where the enthalpy and heat capacity changes for the unfolding of the average globular protein are 6.7 cal/g and 0.12 cal/(K × g), respectively [see Methods and (Robertson and Murphy, 1997)]. Note, the storage conditions of the CoV-2 spike has recently been reported to impact the thermal denaturation profile of the spike suggesting substantial thermal unfolding hysteresis (Edwards et al., 2020).

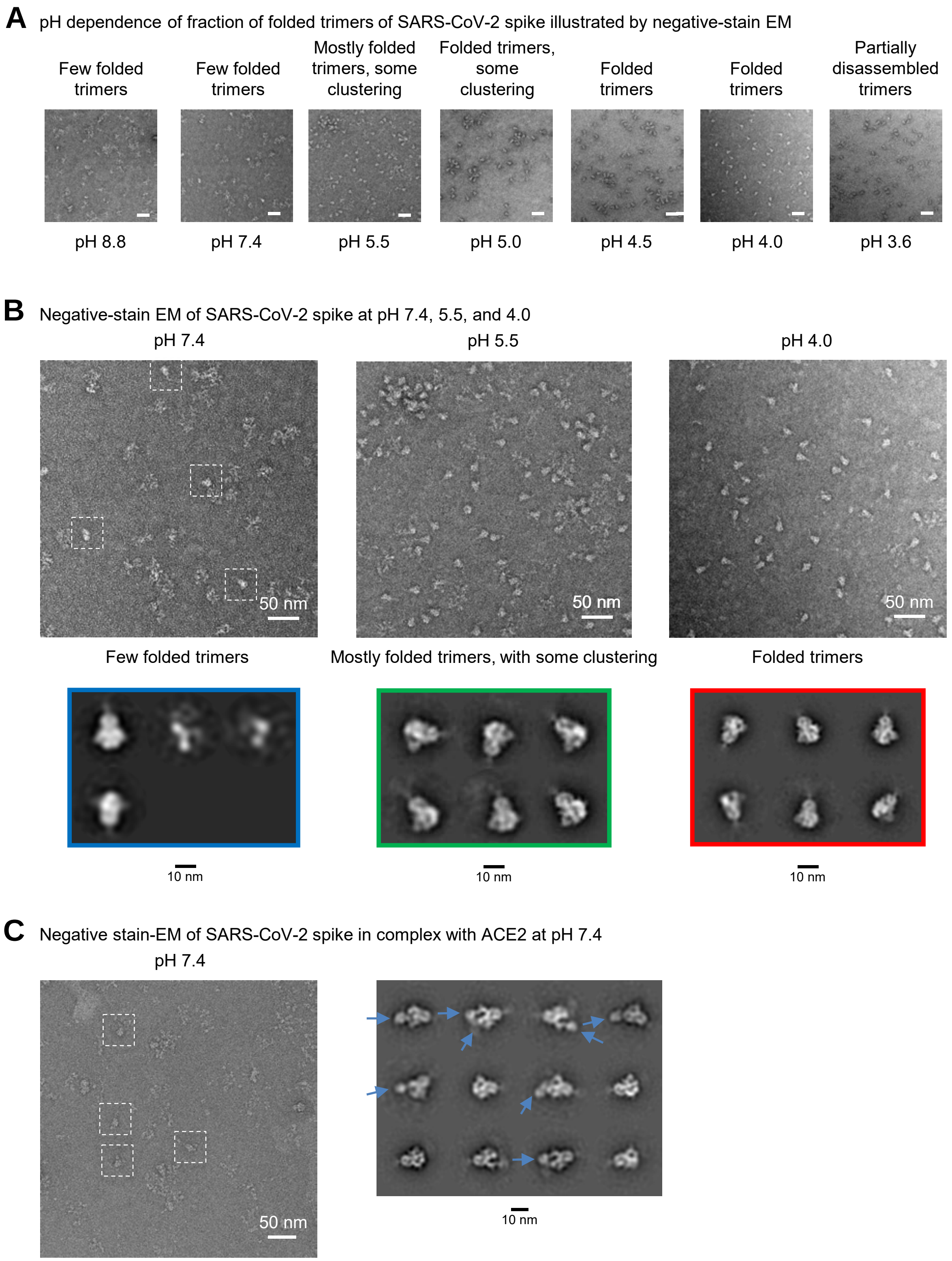


**Figure S2. Negative-Stain EM Indicates a pH Dependence in the Fraction of Folded SARS-CoV-2 Spike, with More Assembled Trimers at Moderately Acidic pH, Related to Figure 1**. (**A**) Typical micrographs in the 3.6-8.8 pH range. The fraction of fully assembled spike trimers is increased at pH 5.5–4.0. The same protein sample was used in all experiments. Scale bars are 50 nm long. (**B**) Close-up views and 2D-class averages for spike at pH 7.4, 5.5, and 4.0. Left: fully assembled spike trimers are present at pH 7.4 (white boxes) but constitute a minor fraction. Center: at pH 5.5, most spikes are folded and fully assembled, with some clustering. Right: at pH 4.0, mostly individual, fully assembled spike trimers are present. Note that the scarcity of folded species at pH 7.4 does not preclude high-resolution cryo-EM analysis provided that a sufficient amount of data is collected. (**C**) Negative-stain EM resolved spike-ACE2 complexes at pH 7.4 despite the presence of a large fraction of spike molecules that did not form folded trimers. Left: a typical micrograph; complexes with folded spikes are boxed. Right: 2D class average images. Arrows point to bound ACE2 monomers.

### Figure S3

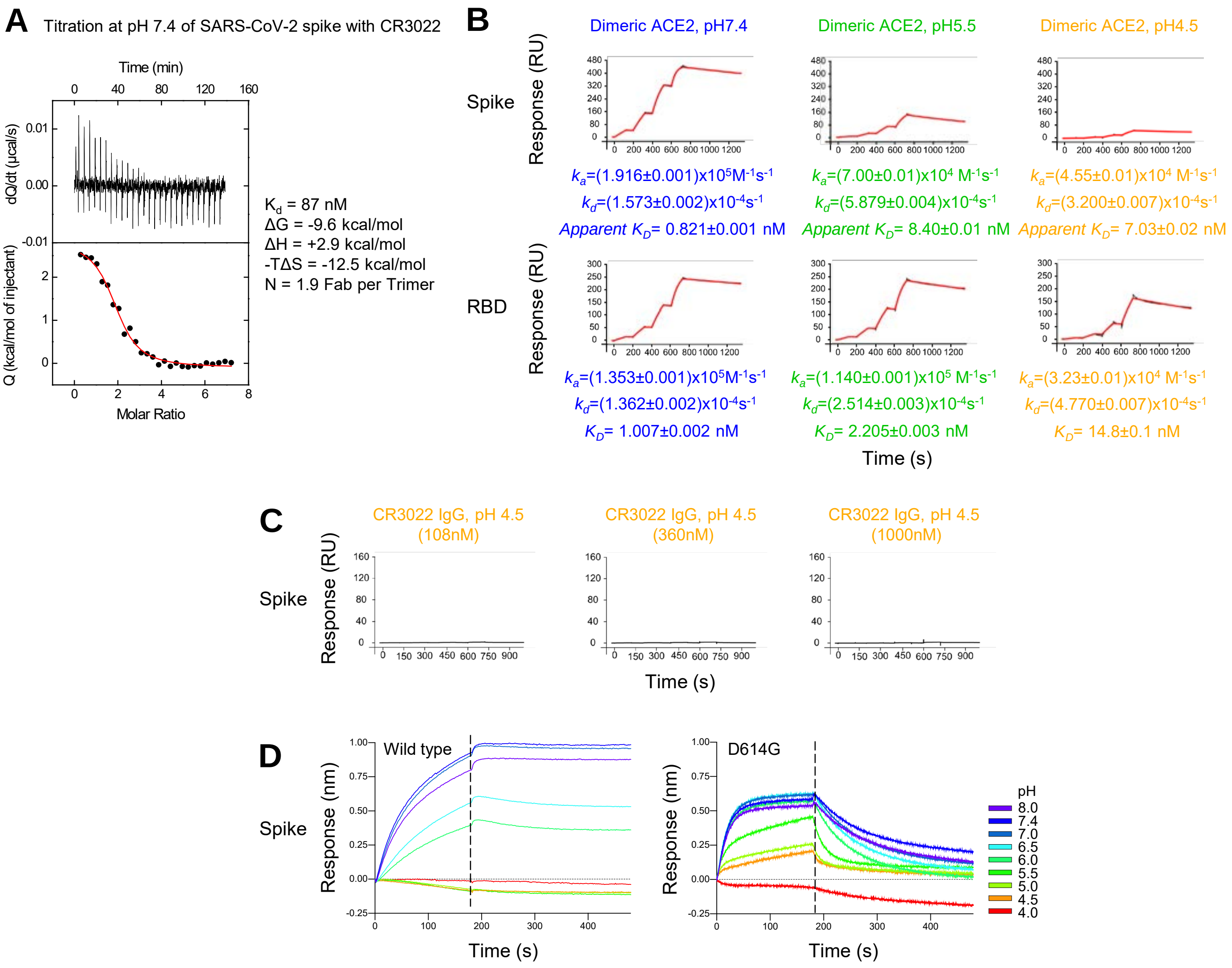


**Figure S3. Isothermal Titration Calorimetry, Surface Plasmon Resonance, and Bio-Layer Interferometry of SARS-CoV-2 Spike Binding to Antibody CR3022 and ACE2 Receptor, Related to Figure 1**. (**A**) Isothermal titration calorimetry of SARS-CoV-2 spike with Fab CR3022 at pH 7.4 and 25 °C. The thermodynamic binding parameters together with the stoichiometry are shown to the right. (**B**) SPR single-cycle kinetics for dimeric ACE2 binding to biotinylated spike (top) and biotinylated-RBD (middle) each tethered to a chip surface. Black traces represent the experimental data and red traces represent the fit to a 1:1 interaction model. The error in each measurement represents the error of the fit. The affinity between trimeric spike and dimeric ACE2 are expressed as apparent $K_D$. (**C**) SPR single cycle kinetics for CR3022 IgG binding to the spike at pH 4.5 performed at higher concentration series, 4.00 to 108 nM (left), 13.3 to 360. nM (middle) and 37.0 to 1000 nM (right). (**D**) Bio-layer interferometry assay of CR3022 IgG binding to SARS-CoV-2 spike and its D614G mutant. Antibody was loaded onto anti-human-Fc sensors and dipped into spike-containing solutions at different pH.

### Figure S4

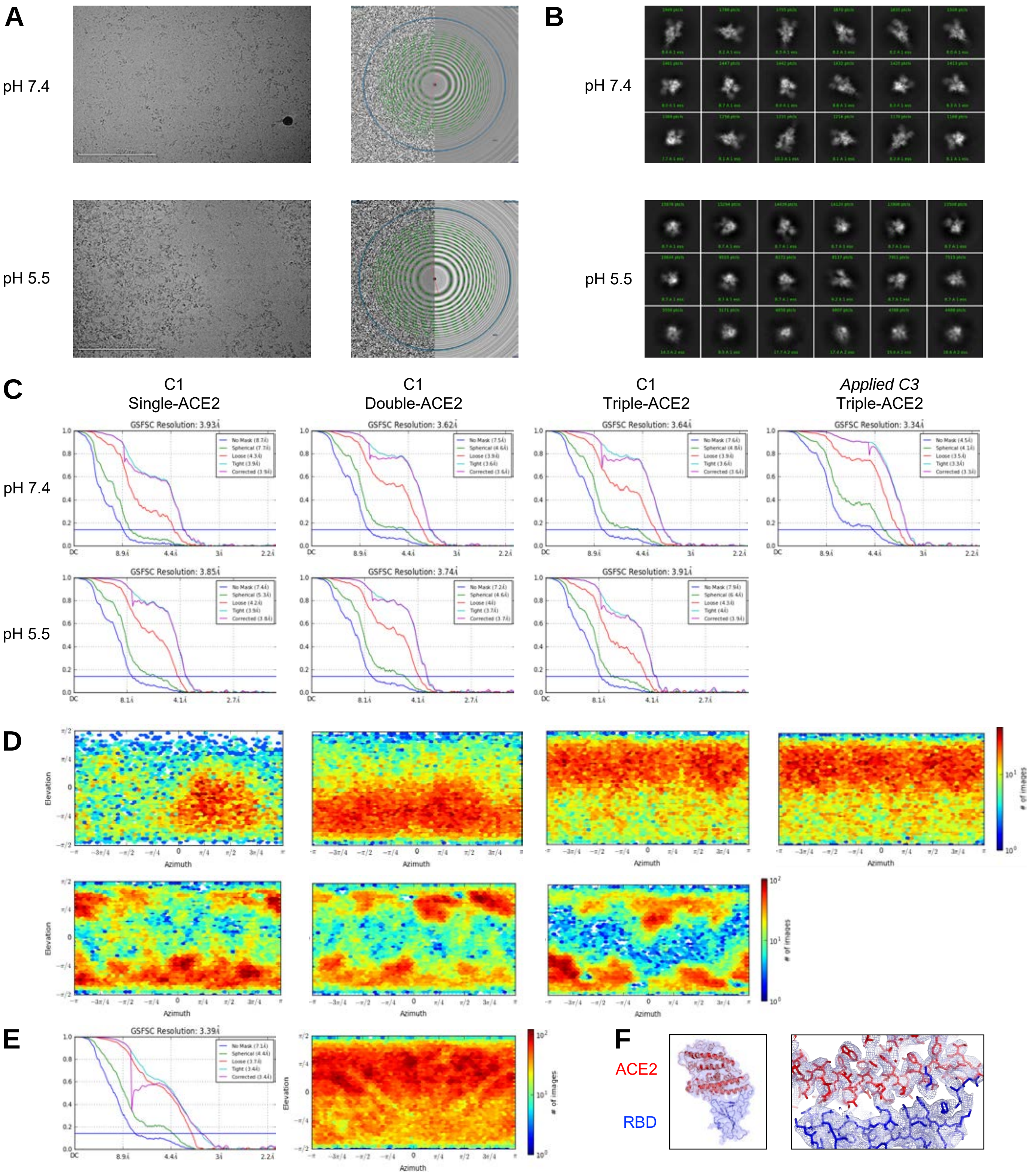


**Figure S4. CryoEM with ACE2 at pH 7.4 and 5.5, Related to Figure 2.** (**A**) Representative micrographs at pH 7.4 (top) and 5.5 (bottom) are shown along with corresponding CTF. (**B**) Representative 2D class averages are shown for each pH. (**C**) The gold-standard Fourier shell correlation is shown for each complex at both pH 7.4 and 5.5 using non-uniform refinement. Applied C3 symmetry on the pseudosymmetric triple-bound complex is shown for pH 7.4. (**D**) The orientations of all particles used in the final refinement are shown as a heatmap. (**E**) Symmetry expansion in C3 for the triple-bound complex was used for a masked local refinement of the RBD+ACE2 region. **(F)** Density is shown for the local refinement of the ACE2-RBD region.

*(continued on next page)*

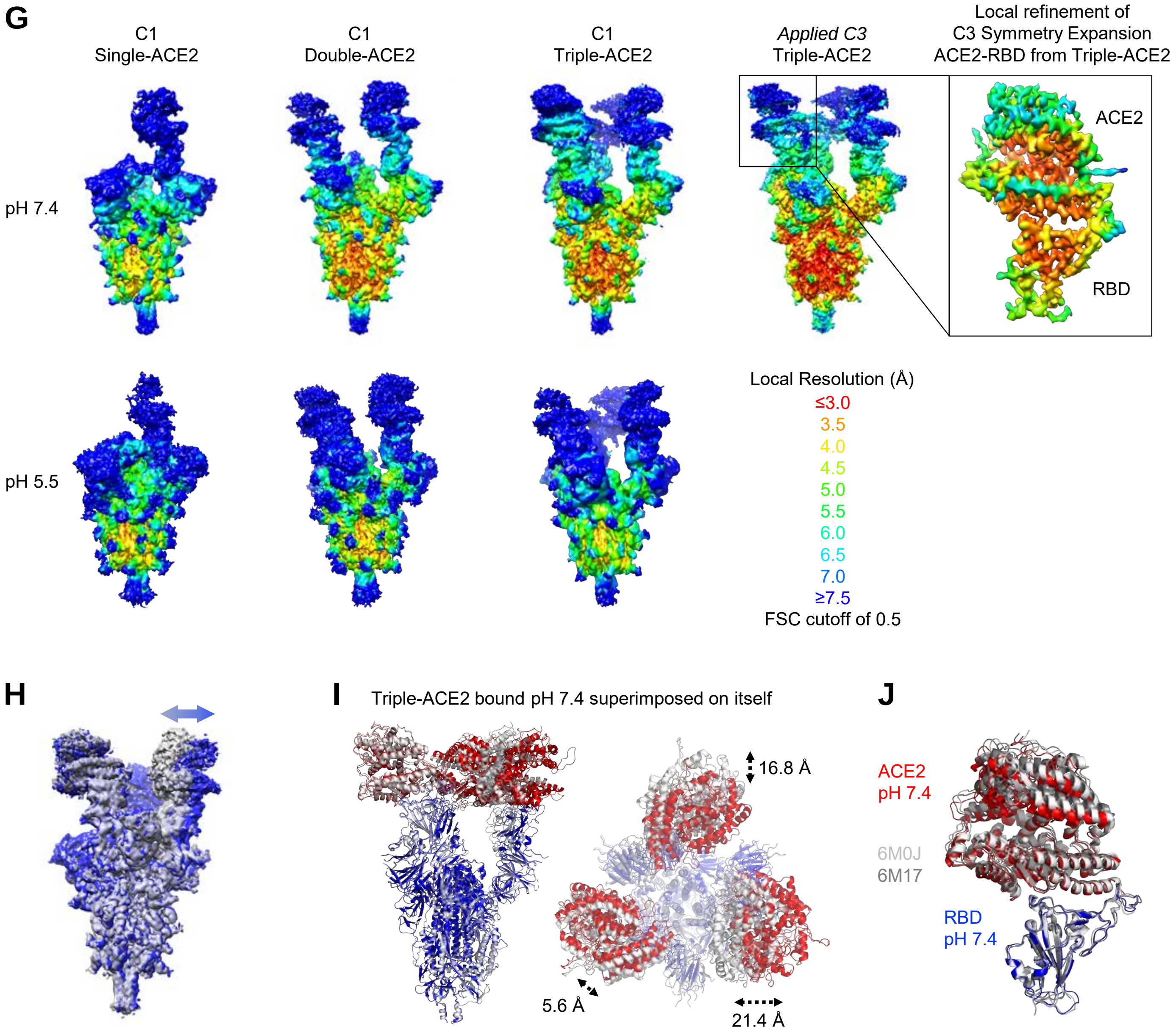


**Figure S4. CryoEM with ACE2 at pH 7.4 and 5.5, Related to Figure 2.** (**G**) The local resolution of each complex is represented. The triple-ACE complex symmetry is broken by the movement of the RBD-ACE2 region. Applied C3 symmetry followed by symmetry expansion and focused refinement results in a well resolved ACE2-RBD region (top right) that was used in all models as a reference structure. (**H**) Triple-bound ACE2 complex map aligned with itself rotated on the 3-fold axis shows the movement of the RDB-ACE2 region. (**I**) **S**tructure of the triple-ACE2 bound pH 7.4 aligned to a copy of itself by S2 shows asymmetry in the RBD-ACE2 angles. (**J**) Comparison of the RBD-ACE2 to PDB IDs 6M0J (light grey) and 6M17 (dark grey) aligned by the RBD shows the ACE2 slightly more closed in these structures. We note that a single ACE2-bound structure of spike was recently described (Xu et al., 2020).

Figure S5

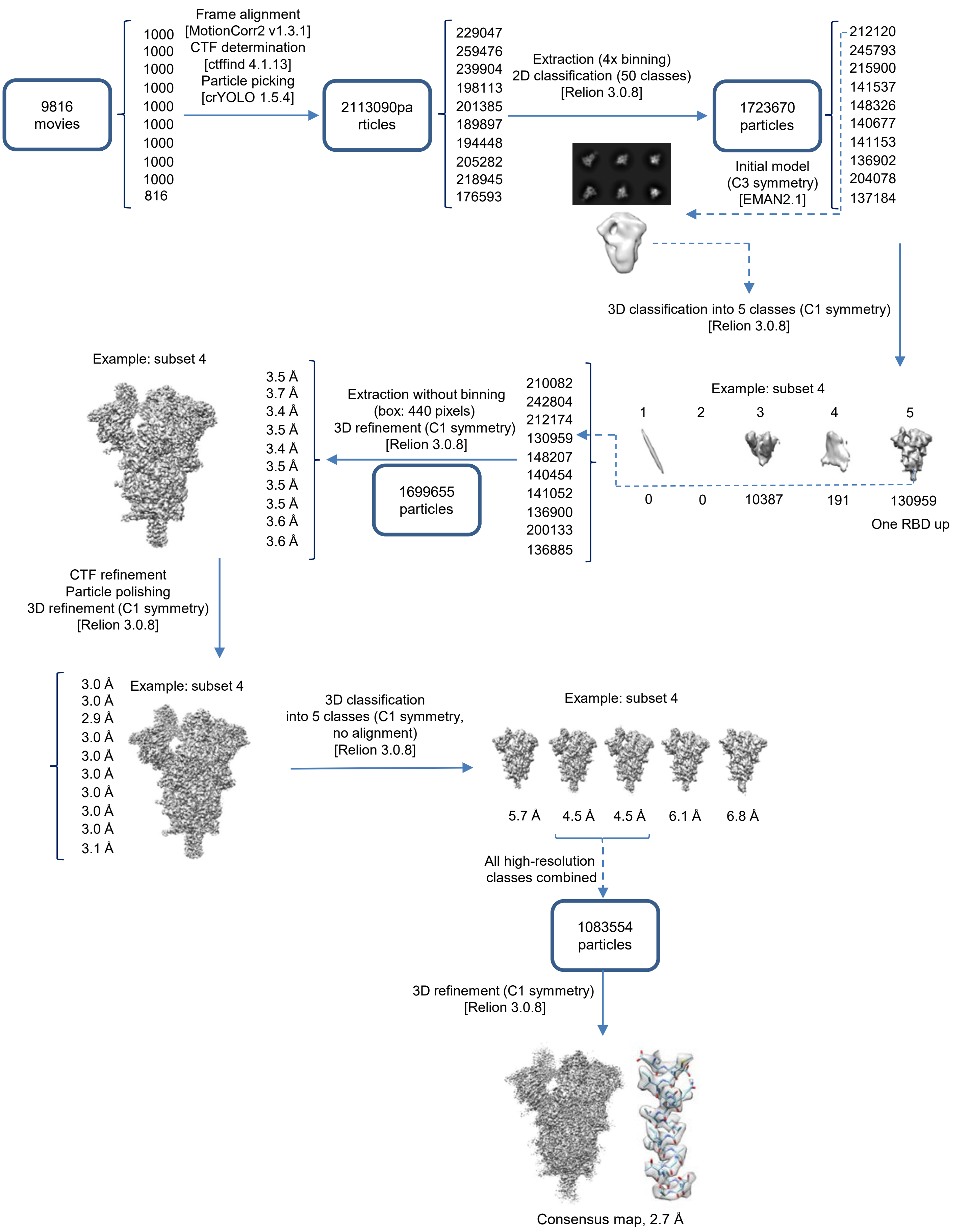


**Figure S5A. Cryo-EM Data Processing Workflow Leading to the Consensus Structure of SARS-CoV-2 Spike at pH 5.5, Related to Figure 3.** Software packages are indicated in square brackets. *(continued on next page)*

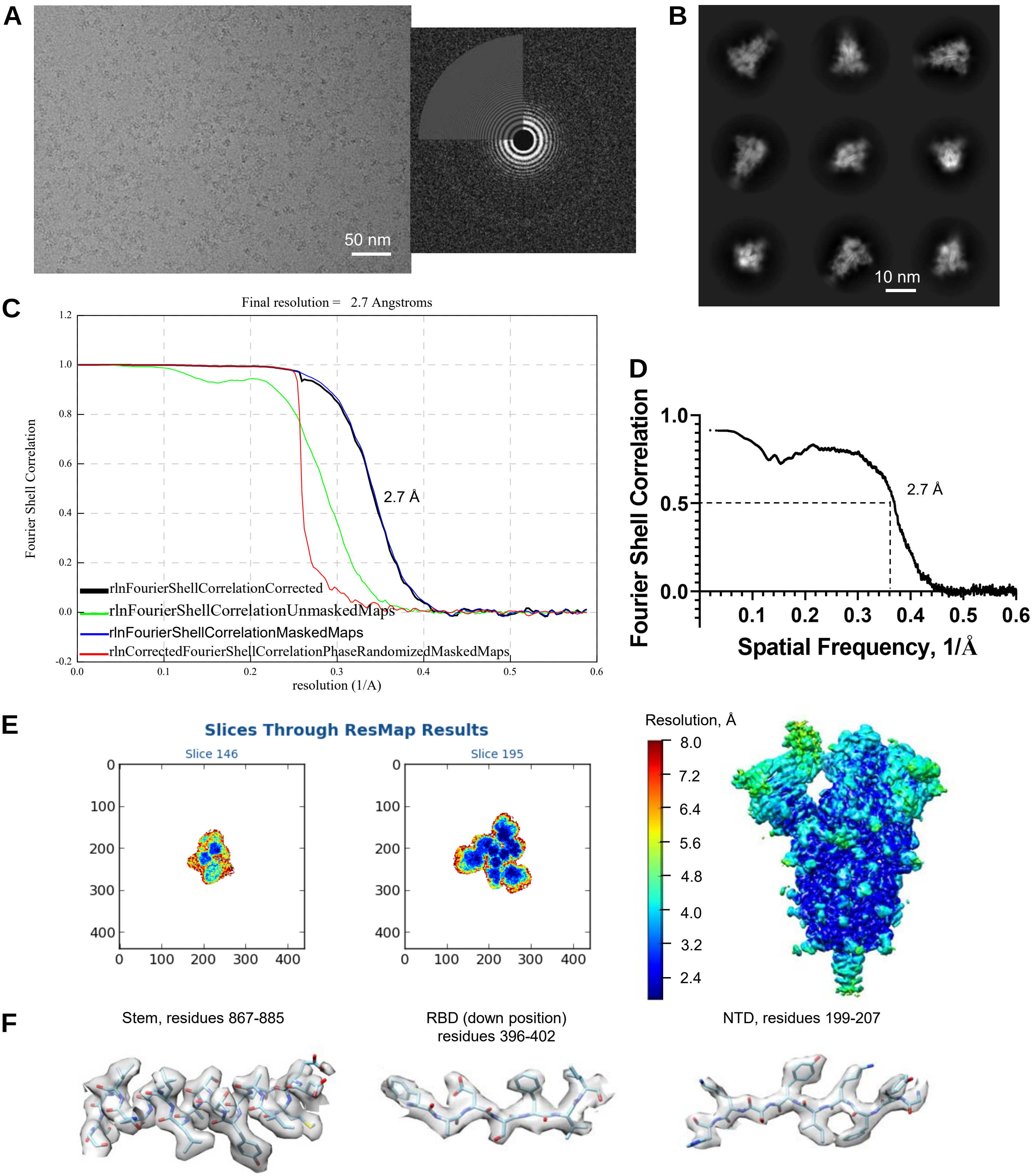


**Figure S5B. Validation of the Consensus CryoEM Map of SARS-CoV-2 Spike at pH 5.5, Related to Figure 3.** (**A**) Representative micrograph (left) and its power spectrum (right). (**B**) Representative high-resolution 2D class averages. (**C**) Gold-standard resolution data generated by Relion. At the 0.143 threshold, the resolution is 2.7 Å. (**D**) Fourier shell correlation curve between the map and the atomic model. (**E**) Results of local resolution analysis using ResMap. Left: Slices through the map at two different levels. Right: Map colored according to local resolution. (**F**) Cryo-EM density in various regions.

*(continued on next page)*

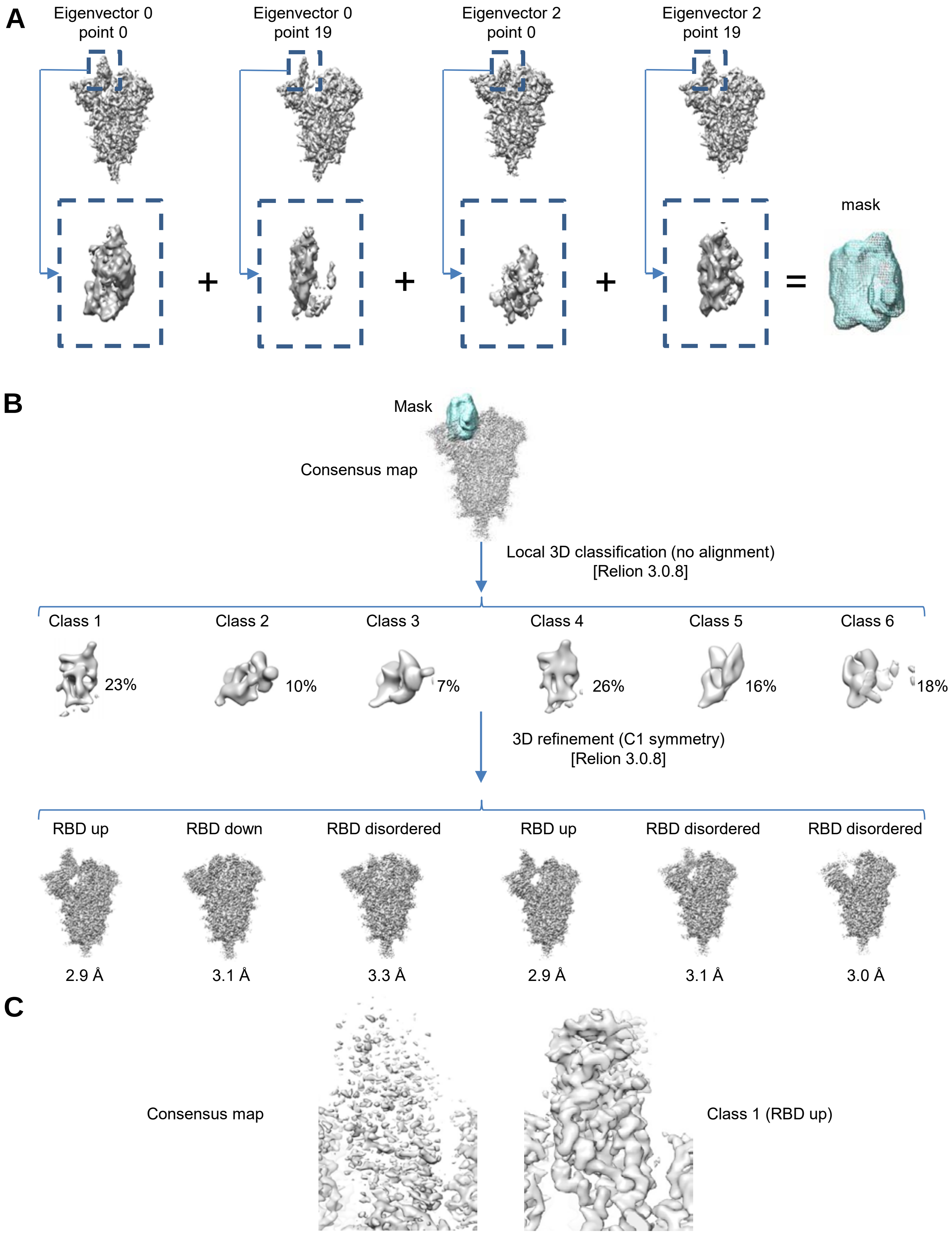


**Figure S5C. Analysis of Heterogeneity in the RBD Positions at pH 5.5, Related to Figure 3.** (**A**) Extreme points (structures) along the trajectories defined by eigenvectors 0 and 2 in 3D variability analysis of the consensus structure were used to create a mask approximating the conformational space of the dynamic RBD domain. (**B**) Local 3D classification of the consensus map within the mask defined in **a** produced six classes. Global 3D refinement of the corresponding subsets of particles resulted in two structures with the RBD in the up position, one with the RBD in the down position, and three with no defined position of the RBD. (**C**) Comparison of the cryo-EM density corresponding to the RBD in the up position between the consensus map and the Class 1 map. The RBD is fully defined in the latter case. *(continued on next page)*

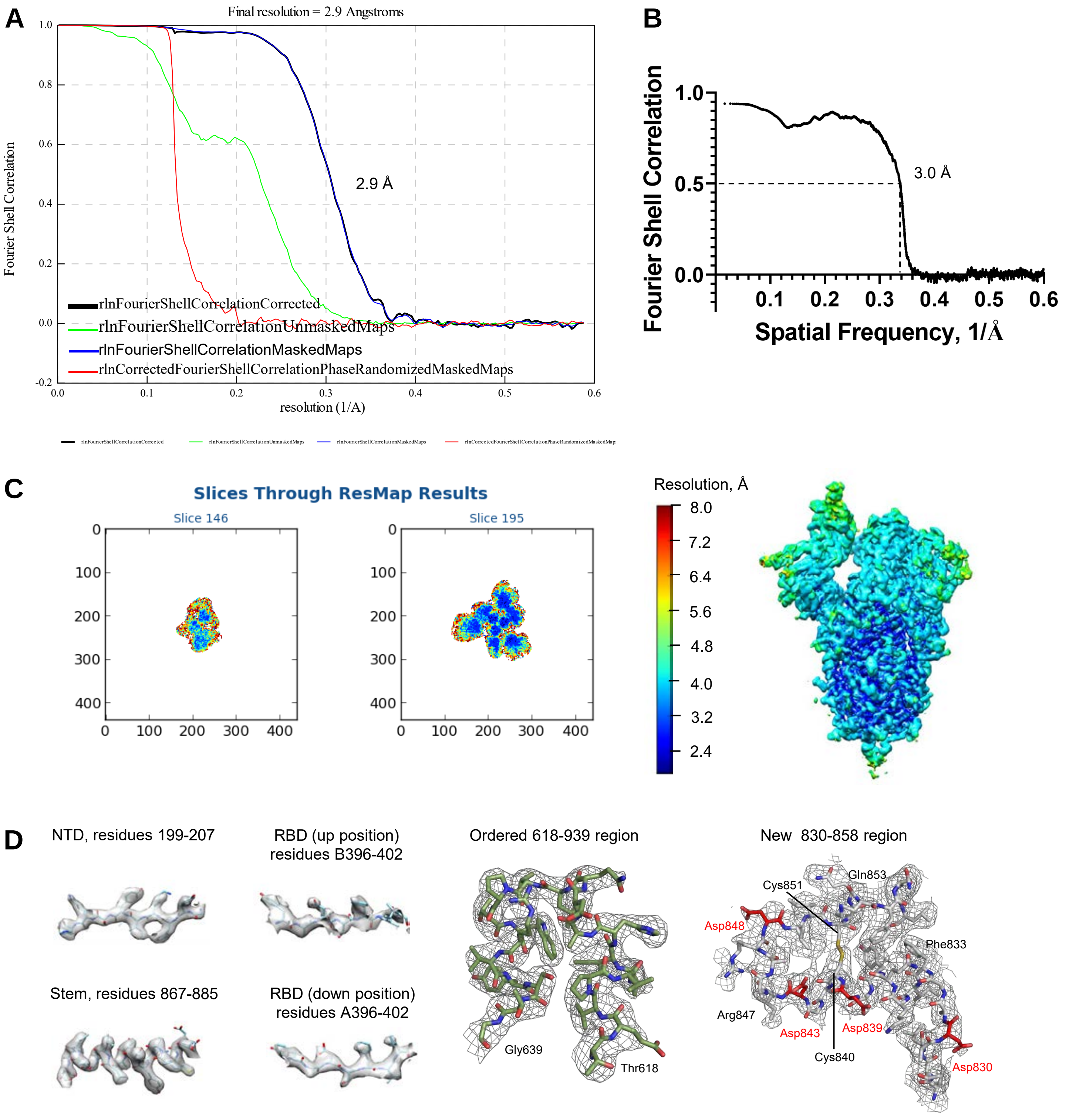


**Figure S5D. Validation of the Cryo-EM Map of SARS-CoV-2 Spike at pH 5.5 with Single RBD in the Up Position (Conformation Up-1), Related to Figure 3.** (**A**) Gold-standard resolution data generated by Relion. At the 0.143 threshold, the resolution is 2.9 Å. (**B**) Fourier shell correlation curve between the map and the atomic model. (**C**) Results of local resolution analysis using ResMap. Left: Slices through the map at two different levels. Right: Map colored according to local resolution. The RBD in the up position is fully resolved. (**D**) Examples of cryo-EM density in various regions. The quality of the density allowed building of atomic model for the RBD in the up position, the 618-639 region and a new conformation of the 830-858 region.

*(continued on next page)*

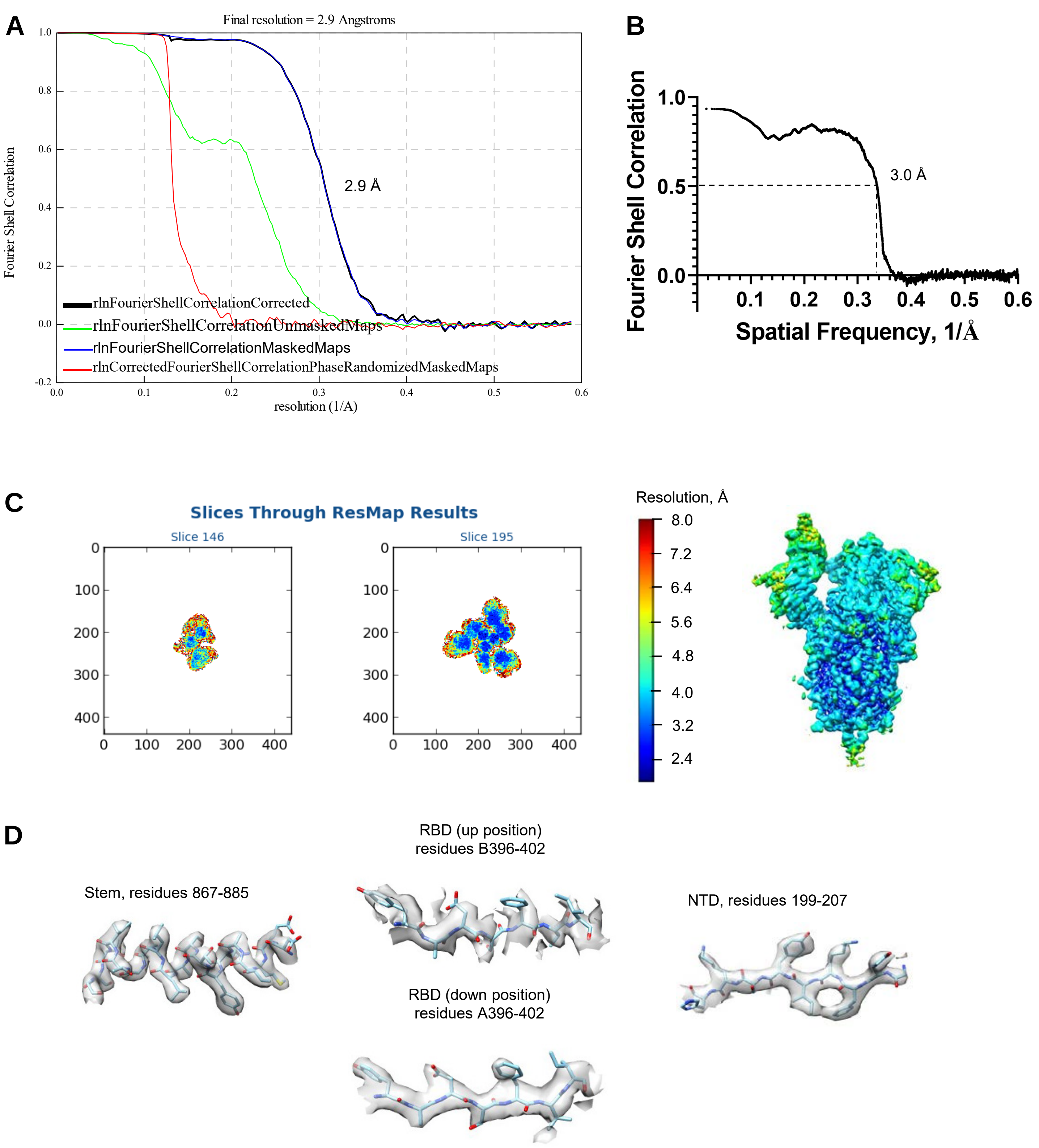


**Figure S5E. Validation of the Cryo-EM Map of SARS-CoV-2 Spike at pH 5.5 with Single RBD in the Up Position (Conformation Up-2), Related to Figure 3.** (**A**) Gold-standard resolution data generated by Relion. At the 0.143 threshold, the resolution is 2.9 Å. (**B**) Fourier shell correlation curve between the map and the atomic model. (**C**) Results of local resolution analysis using ResMap. Left: Slices through the map at two different levels. Right: Map colored according to local resolution. The RBD in the up position is fully resolved. (**D**) Examples of cryo-EM density in various regions. The quality of the density for the RBD in the up position allowed building an atomic model for this domain. *(continued on next page)*

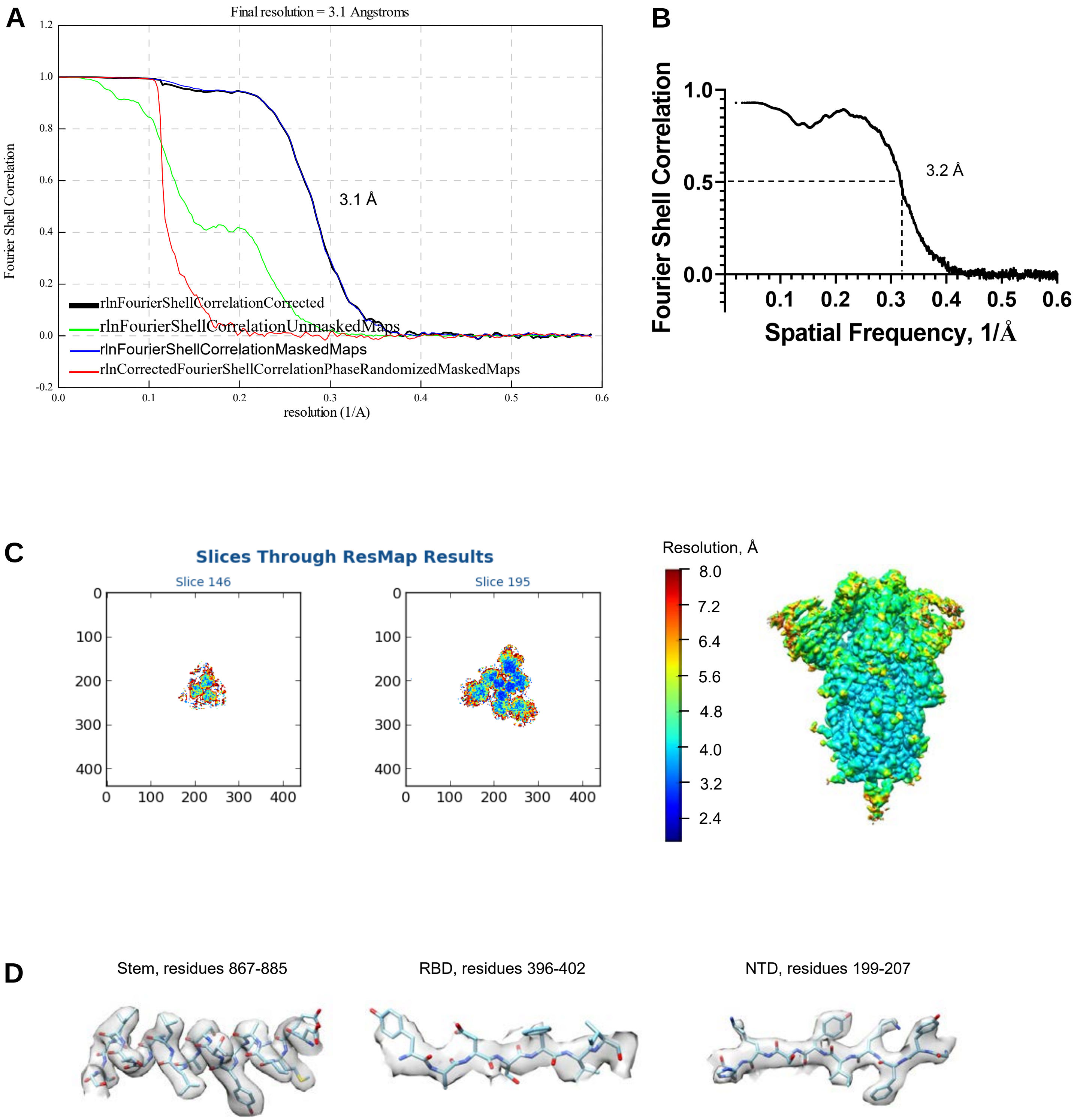


**Figure S5F. Validation of the Cryo-EM Map of SARS-CoV-2 Spike at pH 5.5 with All RBDs in the Down Position, Related to Figure 3.** (**A**) Gold-standard resolution data generated by Relion. At the 0.143 threshold, the resolution is 3.1 Å. (**B**) Fourier shell correlation curve between the map and the atomic model. (**C**) Results of local resolution analysis using ResMap. Left: Slices through the map at two different levels. Right: Map colored according to local resolution. (**D**) Examples of cryo-EM density in various regions.

*(continued on next page)*

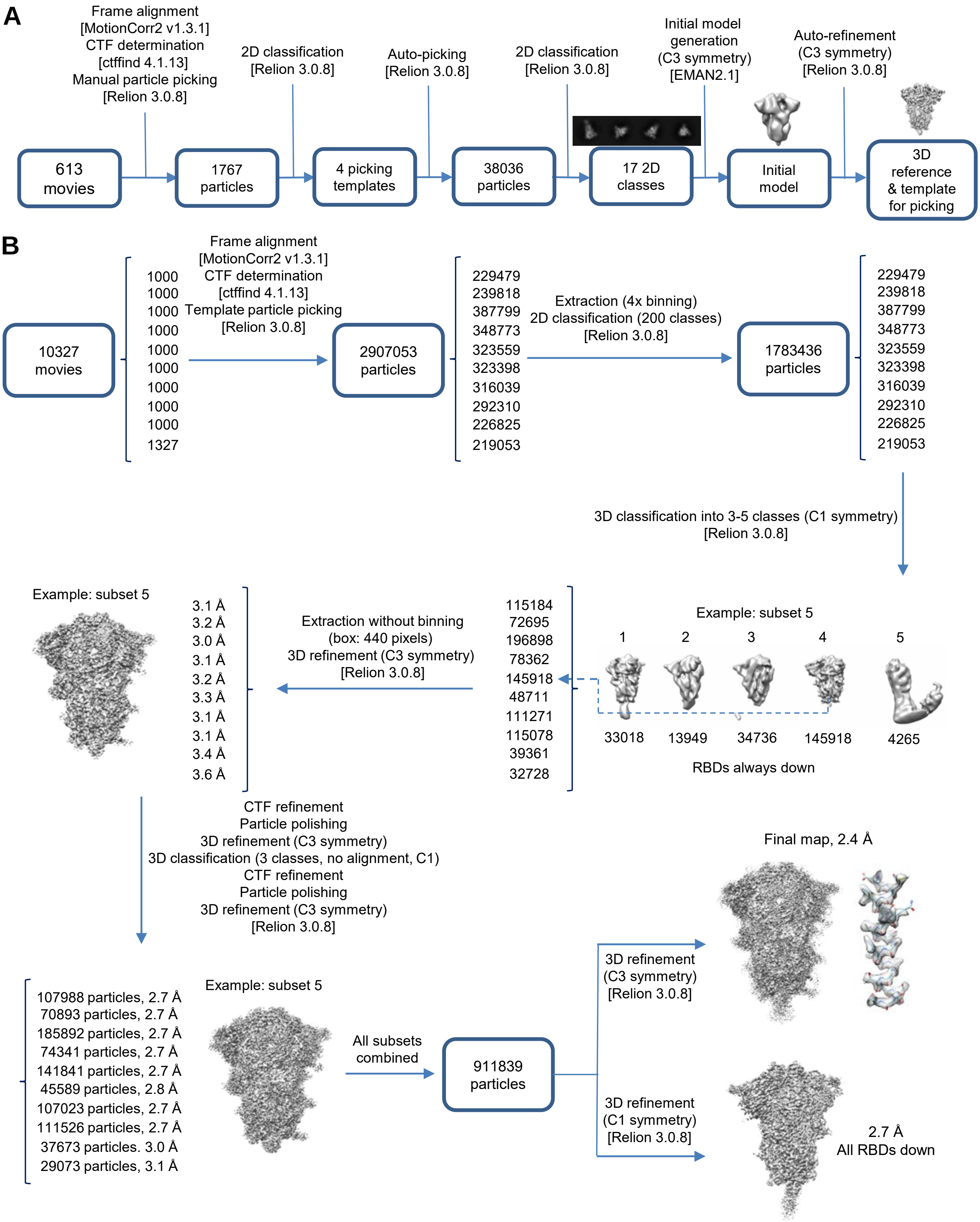


**Figure S5G. Cryo-EM Data Processing Workflow for SARS-CoV-2 Spike at pH 4.0, Related to Figure 3.** (**A**) generation of the initial 3D reference (also used as the 3D particle picking template). (**B**) Steps of single particle analysis. Software packages are indicated in square brackets. *(continued on next page)*

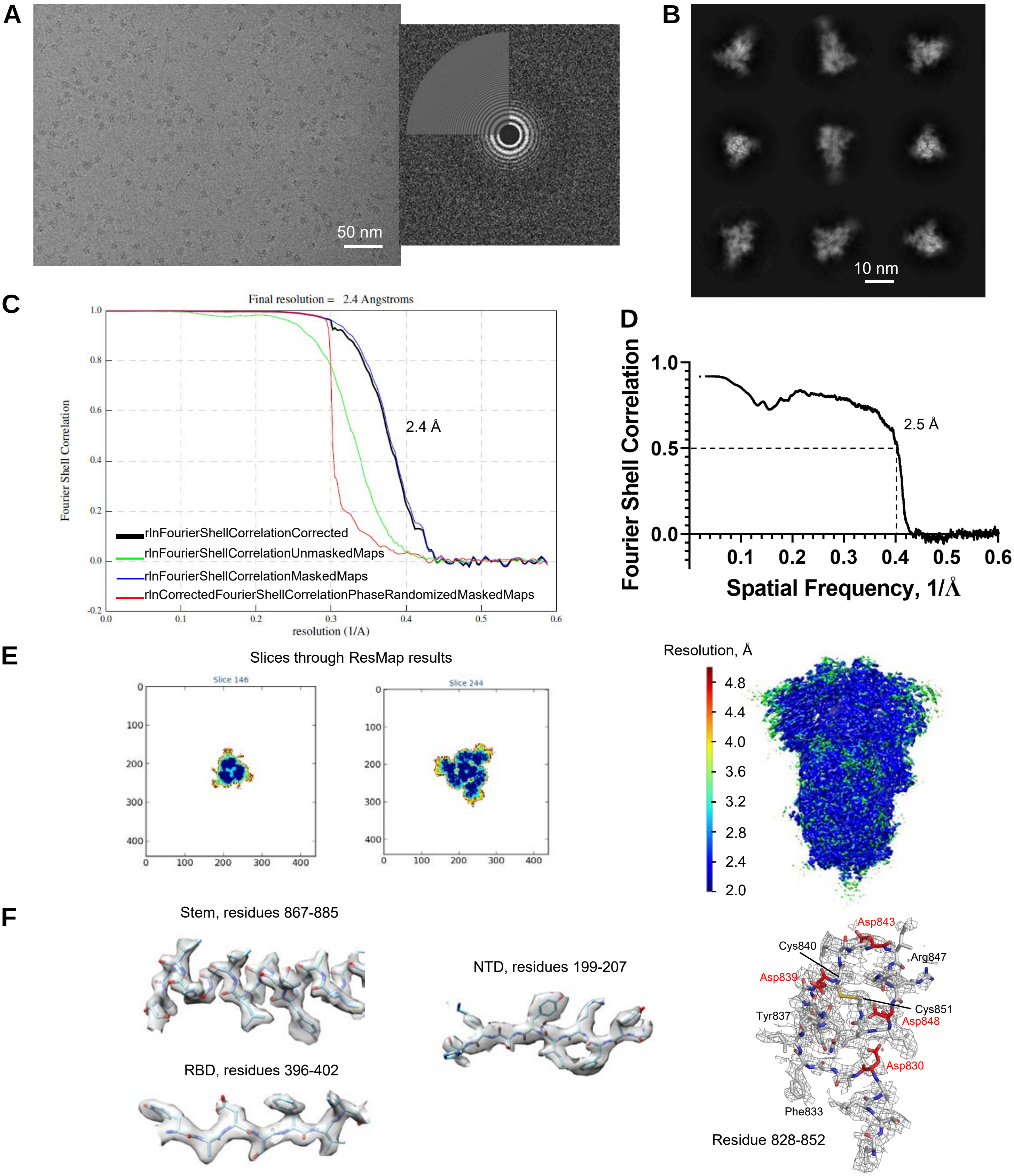


**Figure S5H. Validation of the Cryo-EM Map of SARS-CoV-2 Spike at pH 4.0 Refined with C3 Symmetry Imposed, Related to Figure 3.** (**A**) Representative micrograph (left) and its power spectrum (right). (**B**) Representative high-resolution 2D class averages. (**C**) Gold-standard resolution data generated by Relion. At the 0.143 threshold, the resolution is 2.4 Å. (**D**) Fourier shell correlation curve between the map and the atomic model. (**E**) Results of local resolution analysis using ResMap. Left: Slices through the map at two different levels. Right: Map colored according to local resolution. (**F**) Examples of cryo-EM density in various regions.

*(continued on next page)*

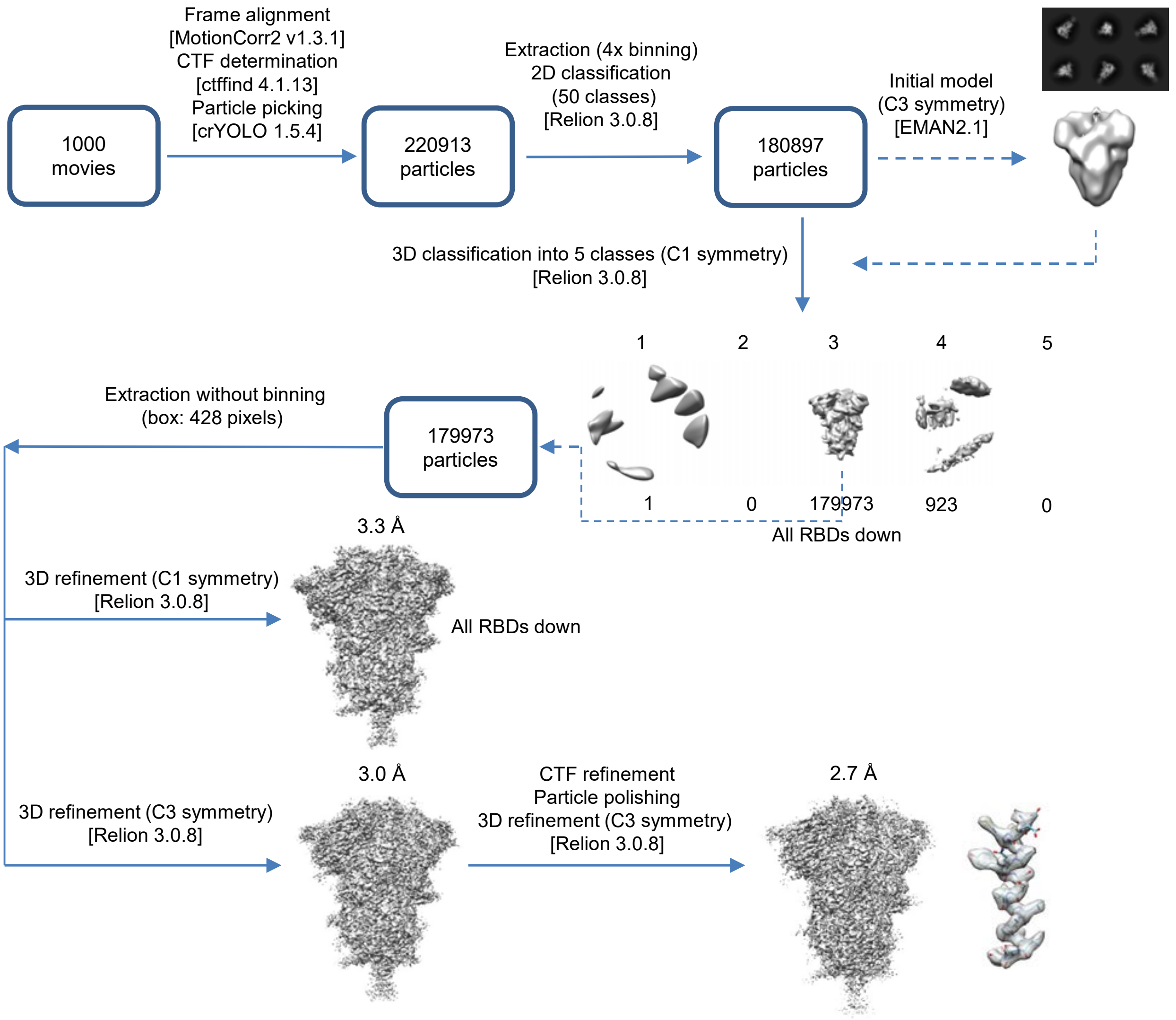


**Figure S5I. Cryo-EM Data Processing Workflow Leading to the Structure of SARS-CoV-2 S at pH 4.5. Related to Figure 3.** Software packages are indicated in square brackets. RBD: receptor binding domain. *(continued on next page)*

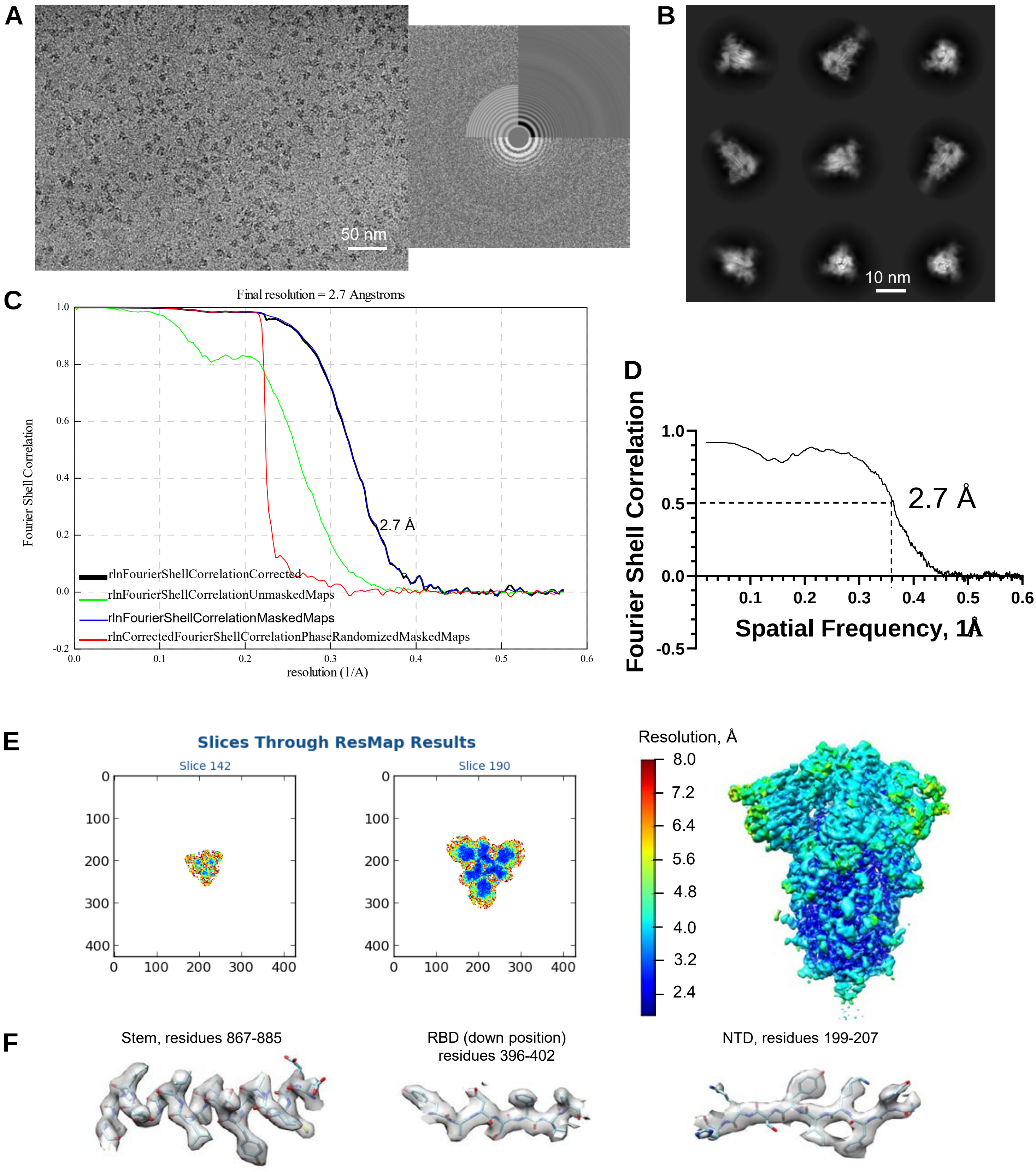


**Figure S5J. Validation of the CryoEM Map of SARS-CoV-2 S at pH 4.5. Related to Figure 3.** (**A**) Representative micrograph (left) and its power spectrum (right). (**B**) Representative high-resolution 2D class averages. (**C**) Gold-standard resolution data generated by Relion. At the 0.143 threshold, the resolution is 2.7 Å. (**D**) Fourier shell correlation curve between the map and the atomic model. (**E**) Results of local resolution analysis using ResMap. Left: Slices through the map at two different levels. Right: Map colored according to local resolution. (**F**) Examples of cryo-EM density in various regions.

**A**

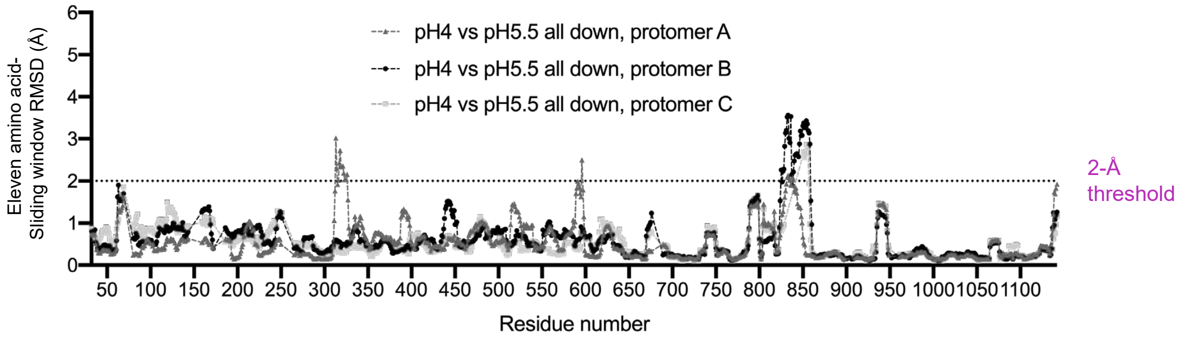


**B**

| Refolding regions with >2-Å RMSD | | | 313-326 | 523-530 | 824-858 |
|---|---|---|---|---|---|
| pH 5.5 single up versus pH4 | Protomer A | 2-Å threshold | 313-325 | | |
| | | (mean, max RMSD) | (2.33, 3.06) | (0.93, 1.36) | (0.59, 1.21) |
| | | Disordered | | | 829-847 |
| | Protomer B | 2-Å threshold | | 523-530 | 829-858 |
| | | (mean, max RMSD) | (0.49, 0.78) | (3.04, 3.84) | (2.73, 3.50) |
| | | Disordered | | | |
| | Protomer C | 2-Å threshold | | (0.42, 0.48) | 824-841, 851-858 |
| | | (mean, max RMSD) | (0.36, 0.53) | | (2.82, 3.84) |
| | | Disordered | | | 842-850 |
| pH 5.5 all down versus pH 4 | Protomer A | 2-Å threshold | 313-326 | | 832-835, 838-839 |
| | | (mean, max RMSD) | (2.28, 3.02) | (0.97, 1.26) | (1.29, 2.14) |
| | | Disordered | | | |
| | Protomer B | 2-Å threshold | | | 829-858 |
| | | (mean, max RMSD) | (0.52, 0.60) | (0.70, 0.73) | (2.84, 3.56) |
| | | Disordered | | | |
| | Protomer C | 2-Å threshold | | | 851-856 |
| | | (mean, max RMSD) | (0.33, 0.44) | (0.42, 0.55) | (1.81, 2.86) |
| | | Disordered | | | 829-849 |

**C**

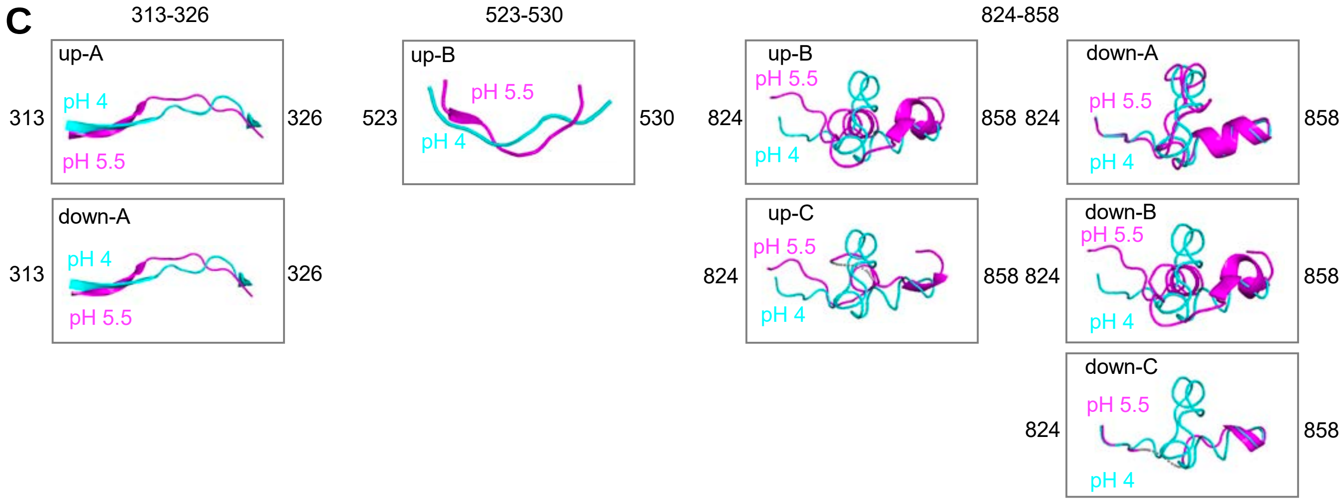


**Figure S6. Details of Refolding Region Analysis, Related to Figures 4 and 5**. (**A**) Identification of refolding regions through rmsd analysis with a 11-residue sliding window comparing pH 5.5 all-down and 4.0 structures, with rmsds calculated for backbone atoms. (**B**) Statistics on refolding analysis comparing pH 4.0 structure to pH 5.5 single-up and pH 5.5 all-down structures respectively, including regions with greater than 2-Å rmsd, mean and maximum rmsd, and disordered residues in pH 5.5 structures, for three refolding regions. (**C**) Alignment of refolding regions between comparing pH 4.0 structure and pH 5.5 structures. pH 4.0 structures are colored in cyan. pH 5.5 structures are colored in magenta. Missing residues are denoted with gray dash lines.

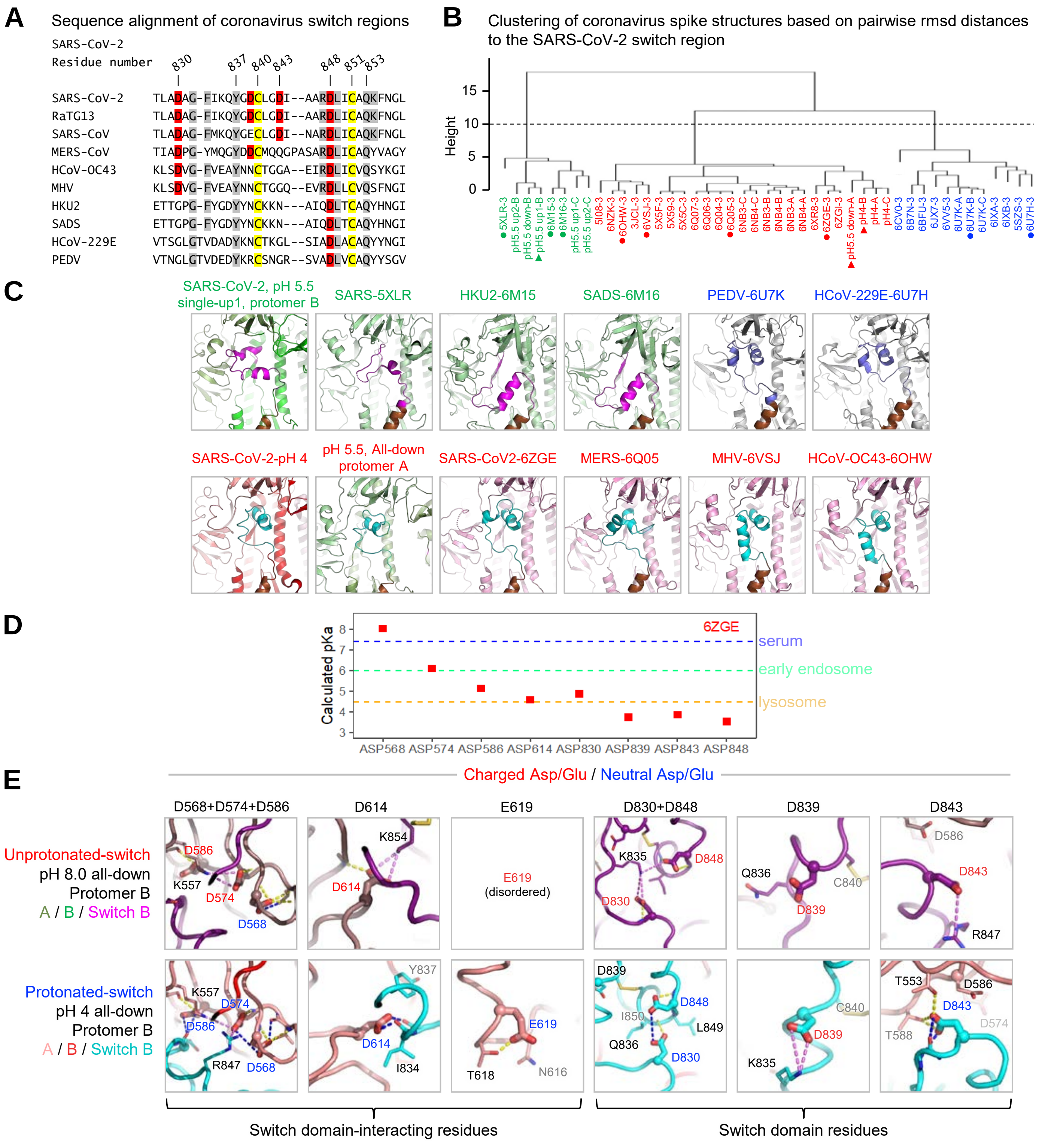


**Figure S7. Structural Comparison of the Switch Regions in Different Coronaviruses, Related to Figures 4-6.** (**A**) Alignment of the switch region sequences from various coronavirus. (**B**) Hierarchical clustering of coronavirus spike structures based on pairwise rmsd distances of the switch region (residues 824-858, SARS-CoV-2 numbering). Only structures with at least 70% of the residues (25 residues) defined for the switch region were considered in the analysis. Entries were labeled as PDB ID-Chain (except for the structures in the current paper). Only one chain per trimer was included for structures with C3 symmetry. Three main clusters were identified, and the members of each cluster were colored green, red, and blue, respectively. ▲ and ● indicate entries for comparison in (A) and (C). (**C**) Conformation of the switch regions. Representative structures are shown for each of the 3 clusters identified in (B). Switches with similar conformation to that in protomer B of the pH 5.5 single-up conformation are shown in magenta. Switches with similar conformation to that in the pH 4.0 structure are shown in cyan. Switches in the third cluster are shown in light blue.

*(figure legend continued on next page)*

**Figure S7.** (continued from the previous page) (**D**) PROPKA-calculated pKas for pH-dependent switch region Asp residues in the recent pH 8.0 all-down SARS-CoV-2 spike structure (PDB 6zge). All three chains have the same pKa for each residue due to imposed C3 symmetry. Typical pH values for serum (7.4), early endosome (6.0) and late endosome (4.5) are indicated by dashed lines, each colored as in Figures 1A and 6. The calculated pKa values indicate that only Asp568 will likely be protonated for a switch region in this conformation at pH 8.0. (**E**) Close-up views of Asp residues from (D) in comparison with those in the pH 4.0 structure (Figure 6) reveal changes in chemical environment for each residue. Highlighted residue labels are colored based on predicted protonation state with charged Asp/Glu in red, and neutral (protonated) Asp/Glu in blue. Side chains of the highlighted residues are shown as thick sticks, those of nearby residues within 4 Å shown as thin sticks. Dashed lines indicate hydrogen bonds (yellow) and salt bridges (violet), with hydrogen bonds that require Asp protonation indicated in blue. The pKa shifts between unprotonated- and protonated-switch conformations define a pH-dependent stability gradient that favors the protonated-switch form at lower pH (Yang & Honig, 1993). However, other factors such as global conformational constraints may also play a role in favoring one conformation over another. Indeed, 6xr8 (Cai et al. 2020) and 6zge (Wrobel et al., 2020) are part of a number of recently reported SARS-CoV-2 spike structures including (Xiong et al., 2020; Xu et al., 2020) that contain switch domain conformations at serological or basic pH that are similar to, yet clearly distinct from, the switch conformations we report at lower pH.

### Table S1. Cryo-EM Data Collection, Refinement and Validation Statistics for ACE2-Bound Structures at pH 7.4 and 5.5, Related to Figure 2.

**Cryo-EM data collection, refinement and validation statistics**

| | SARS-CoV-2 spike with single ACE2 at pH 7.4 | SARS-CoV-2 spike with double ACE2 at pH 7.4 | SARS-CoV-2 spike with triple ACE2 at pH 7.4 | Focused ACE2-RBD at pH 7.4, after C3 Symmetry Expansion | SARS-CoV-2 spike with single ACE2 at pH 5.5 | SARS-CoV-2 spike with double ACE2 at pH 5.5 | SARS-CoV-2 spike with triple ACE2 at pH 5.5 |
|---|---|---|---|---|---|---|---|
| **Data collection and processing** | | | | | | | |
| Magnification | 81,000 | 81,000 | 81,000 | 81,000 | 81,000 | 81,000 | 81,000 |
| Voltage (kV) | 300 | 300 | 300 | 300 | 300 | 300 | 300 |
| Electron exposure (e–/Å$^2$) | 53.49 | 53.49 | 53.49 | 53.49 | 51.3 | 51.30 | 51.3 |
| Defocus range (μm) | -0.4 to -3.6 | -0.4 to -3.6 | -0.4 to -3.6 | -0.4 to -3.6 | -0.2 to -3.7 | -0.2 to -3.7 | -0.2 to -3.7 |
| Pixel size (Å) | 1.058 | 1.058 | 1.058 | 1.058 | 1.058 | 1.058 | 1.058 |
| Symmetry imposed | C1 | C1 | C1 | C1 | C1 | C1 | C1 |
| Final particle images (no.) | 16,997 | 48,008 | 42,947 | 128,841 | 46,714 | 55,297 | 47,386 |
| Map resolution (Å) | 3.93 | 3.62 | 3.64 | 3.39 | 3.85 | 3.74 | 3.91 |
| FSC threshold | 0.143 | 0.143 | 0.143 | 0.143 | 0.143 | 0.143 | 0.143 |
| **Refinement** | | | | | | | |
| Initial model used (PDB code) | 6VXX, 6M0J | 6VXX, 6M0J | 6VXX, 6M0J | 6M0J | 6VXX, 6M0J | 6VXX, 6M0J | 6VXX, 6M0J |
| Model resolution (Å) | 3.9 | 3.7 | 3.7 | 3.4 | 4.1 | 4 | 4.1 |
| FSC threshold | 0.143 | 0.143 | 0.143 | 0.143 | 0.143 | 0.143 | 0.143 |
| Map sharpening *B* factor (Å$^2$) | -58.4 | -74.1 | -61.9 | -77.8 | -60.3 | -55.4 | -51.3 |
| Model composition | | | | | | | |
| Non-hydrogen atoms | 28110 | 32784 | 39741 | 6509 | 27320 | 31342 | 39069 |
| Protein residues | 3589 | 4165 | 4887 | 788 | 3512 | 4123 | 4887 |
| Ligands | 12 | 61 | 71 | 9 | 12 | 61 | 71 |
| *B* factors (Å$^2$)(mean) | | | | | | | |
| Protein | 140.1 | 155.45 | 158.33 | 87.25 | 140.5 | 143.7 | 180.1 |
| Ligand | 182.7 | 141.3 | 182.72 | 99.24 | 143.2 | 168.2 | 198.9 |
| R.m.s. deviations | | | | | | | |
| Bond lengths (Å) | 0.007 | 0.003 | 0.004 | 0.003 | 0.008 | 0.009 | 0.004 |
| Bond angles (°) | 0.843 | 0.697 | 0.0795 | 0.713 | 1.128 | 1.076 | 0.877 |
| Validation | | | | | | | |
| MolProbity score | 1.46 | 1.43 | 1.58 | 1.50 | 2.71 | 2.1 | 1.6 |
| Clash score | 4.44 | 4.3 | 4.63 | 3.70 | 6.42 | 5.42 | 4.93 |
| Poor rotamers (%) | 0.14 | 0 | 0.07 | 0.0 | 0.12 | 0.98 | 0.02 |
| Ramachandran plot | | | | | | | |
| Favored (%) | 94.4 | 96.57 | 94.7 | 95.15 | 92.9 | 94.5 | 95.09 |
| Allowed (%) | 5.4 | 3.43 | 5.28 | 4.85 | 7.1 | 5.5 | 4.85 |
| Disallowed (%) | 0.1 | 0 | 0.02 | 0 | 0 | 0 | 0.06 |

**Table S2. Cryo-EM Data Collection, Refinement and Validation Statistics for Ligand-Free Spike Structures Determined at pH 5.5, Related to Figure 3.**

| Structures | SARS-CoV-2 S pH 5.5 consensus map (EMD-22253) (PDB 6XM0) | SARS-CoV-2 S pH 5.5 RBD up-1 (EMD-22254) (PDB 6XM3) | SARS-CoV-2 S pH 5.5 RBD up-2 (EMD-22255) (PDB 6XM4) | SARS-CoV-2 S pH 5.5 RBD all-down (EMD-22256) (PDB 6XM5) |
|---|---|---|---|---|
| **Data collection and processing** | | | | |
| Magnification | 105,000 | | | |
| Voltage (kV) | 300 | | | |
| Electron exposure (e–/Å$^2$) | 40 | | | |
| Defocus range (μm) | -1.25 to -2.5 | | | |
| Pixel size (Å) | 0.85 | | | |
| Symmetry imposed | C1 | | | |
| Initial particle images (no.) | 2,113,090 | | | |
| Final particle images (no.) | 1,083,554 | 247,605 | 286,302 | 108,053 |
| Map resolution (Å) | 2.7 | 2.9 | 2.9 | 3.1 |
| FSC threshold | 0.143 | | | |
| Map resolution range (Å) | 1.9-8.4 | | | |
| **Refinement** | | | | |
| Initial model used (PDB code) | 6VYB | pH5.5 | pH5.5 | pH 4.0 |
| Model resolution (Å) | 2.7 | 3.0 | 3.0 | 3.1 |
| FSC threshold | 0.5 | 0.5 | 0.5 | 0.5 |
| Map sharpening *B* factor (Å$^2$) | -72.3 | Local sharpening | Local sharpening | -37.9 |
| Model composition | | | | |
| Non-hydrogen atoms | 25201 | 25384 | 25398 | 25393 |
| Protein residues | 3125 | 3143 | 3143 | 3149 |
| Ligands (glycan) | 56 | 60 | 61 | 58 |
| Water | - | - | - | - |
| *B* factors (Å$^2$)(mean) | | | | |
| Protein | 28.3 | 86.0 | 39.6 | 85.5 |
| Ligand | 45.8 | 98.7 | 62.7 | 100.7 |
| Water | | | | |
| R.m.s. deviations | | | | |
| Bond lengths (Å) | 0.010 | 0.012 | 0.010 | 0.008 |
| Bond angles (°) | 1.096 | 1.014 | 1.082 | 0.850 |
| Validation | | | | |
| MolProbity score | 1.50 | 1.60 | 1.54 | 1.61 |
| Clash score | 2.17 | 3.40 | 3.01 | 3.77 |
| Poor rotamers (%) | 0.37 | 0.47 | 0.40 | 0.51 |
| Ramachandran plot | | | | |
| Favored (%) | 91.4 | 92.6 | 93.1 | 93.2 |
| Allowed (%) | 8.4 | 7.3 | 6.8 | 6.8 |
| Disallowed (%) | 0.2 | 0.1 | 0.1 | 0 |

**Table S3. Cryo-EM Data Collection, Refinement and Validation Statistics for the Ligand-Free Spike Structure Determined at pH 4.5 and 4.0, Related to Figure 3.**

| **Structure** | SARS-CoV-2 Spike at pH 4.5 (EMD-xxxx) (PDB xxx) | SARS-CoV-2 Spike at pH 4.0 (EMD-22251) (PDB 6XLU) |
|---|---|---|
| **Data collection and processing** | | |
| Magnification | 105,000 | 105,000 |
| Voltage (kV) | 300 | 300 |
| Electron exposure (e–/Å$^2$) | 40 | 40 |
| Defocus range (μm) | -1 to -2.5 | -1.25 to -2.5 |
| Pixel size (Å) | 0.873 | 0.85 |
| Symmetry imposed | C3 | C3 |
| Initial particle images (no.) | 220,913 | 2,907,053 |
| Final particle images (no.) | 179,973 | 911,839 |
| Map resolution (Å) | 2.7 | 2.4 |
| FSC threshold | 0.143 | 0.143 |
| Map resolution range (Å) | 1.8-5.6 | 1.8-4.8 |
| **Refinement** | | |
| Initial model used (PDB code) | 6XLU | 6VXX |
| Model resolution (Å) | 2.7 | 2.4 |
| FSC threshold | 0.5 | 0.5 |
| Map sharpening *B* factor (Å$^2$) | -47.8 | -61.0 |
| Model composition | | |
| Non-hydrogen atoms | 25860 | 25990 |
| Protein residues | 3178 | 3178 |
| Ligands | NAG: 64 | NAG: 62 |
| Water | 119 | 250 |
| *B* factors (Å$^2$) (mean) | | |
| Protein | 59.7 | 44.7 |
| Ligand | 88.0 | 67.2 |
| Water | 23.7 | 20.2 |
| R.m.s. deviations | | |
| Bond lengths (Å) | 0.009 | 0.011 |
| Bond angles (°) | 1.016 | 1.190 |
| Validation | | |
| MolProbity score | 1.35 | 1.35 |
| Clash score | 2.25 | 1.77 |
| Poor rotamers (%) | 0.07 | 0.40 |
| Ramachandran plot | | |
| Favored (%) | 95.1 | 93.7 |
| Allowed (%) | 4.9 | 6.2 |
| Disallowed (%) | 0 | 0.1 |

**Table S4. Domain Movements Between Single-RBD Up (pH 5.5) and All-RBD Down (pH 4.0) Structures, Related to Figure 4.**

| | | | pH5.5 | | pH 5.5 single-up conformation 1 | | pH 5.5 single-up conformation 2 | | 6vxx | | 6vyb | |
|---|---|---|---|---|---|---|---|---|---|---|---|---|
| | | | All-down | | B-up | | B-up | | All-down | | B-up | |
| | | | Angle (°) * | Displacement (Å) | Angle (°) | Displacement (Å) | Angle (°) | Displacement (Å) | Angle (°) | Displacement (Å) | Angle (°) | Displacement (Å) |
| pH4 | A | NTD | 0.8 | 0.4 | 0.3 | 0.6 | 1.1 | 0.5 | 6.2 | 6.2 | 6.5 | 6.2 |
| | | RBD | 1.2 | 0.6 | 2.4 | 1.3 | 2.7 | 1.4 | 4.6 | 5.6 | 6.4 | 5.3 |
| | | SD1 | 16.4 | 2.8 | 16.7 | 2.8 | 16.7 | 2.8 | 3.5 | 3.1 | 4.2 | 2.8 |
| | | SD2 | 1.3 | 0.2 | 1.2 | 0.2 | 1.2 | 0.3 | 4.3 | 4.7 | 3.2 | 4.7 |
| | | S2 | 0.2 | 2.9 | 0.3 | 1.3 | 0.3 | 1.3 | 0.1 | 1.8 | 0.2 | 1.9 |
| | B | NTD | 11.9 | 8.2 | 11.9 | 8.8 | 11.7 | 8.6 | 7.2 | 6.3 | 8.0 | 7.8 |
| | | RBD | 3.2 | 1.0 | 64.9 | 22.8 | 67.0 | 25.5 | 4.7 | 4.6 | 66.9 | 22.3 |
| | | SD1 | 7.9 | 2.8 | 12.5 | 4.4 | 14.3 | 4.9 | 3.4 | 3.0 | 12.1 | 5.9 |
| | | SD2 | 8.3 | 2.1 | 9.1 | 2.8 | 9.9 | 2.7 | 4.3 | 4.7 | 5.7 | 4.8 |
| | | S2 | 0.4 | 2.7 | 0.3 | 0.2 | 0.2 | 0.2 | 0.1 | 2.1 | 0.1 | 2.0 |
| | C | NTD | 7.9 | 5.1 | 10.7 | 5.1 | 11.5 | 6.4 | 7.3 | 6.3 | 1.2 | 8.1 |
| | | RBD | 0.6 | 0.4 | 1.2 | 0.2 | 1.7 | 0.6 | 4.5 | 5.7 | 3.4 | 6.8 |
| | | SD1 | 0.4 | 0.5 | 1.1 | 0.9 | 1.0 | 1.1 | 3.5 | 3.0 | 2.6 | 3.1 |
| | | SD2 | 4.7 | 0.9 | 7.0 | 0.8 | 8.5 | 1.0 | 4.3 | 4.8 | 7.6 | 4.6 |
| | | S2 | 0.1 | 1.7 | 0.2 | 1.8 | 0.2 | 0.8 | 0.1 | 2.1 | 0.1 | 2.5 |
| pH5.5-1 | A | NTD | 0.6 | 0.1 | | | 0.5 | 0.3 | | | 5.7 | 6.0 |
| | | RBD | 1.5 | 0.8 | | | 0.9 | 0.2 | | | 5.7 | 4.8 |
| | | SD1 | 1.7 | 0.0 | | | 0.8 | 0.1 | | | 11.2 | 3.5 |
| | | SD2 | 0.1 | 0.1 | | | 0.4 | 0.1 | | | 3.1 | 4.6 |
| | | S2 | 0.1 | 4.0 | | | 0.0 | 0.0 | | | 0.1 | 0.8 |
| | B-up | NTD | 1.3 | 0.6 | | | 0.6 | 0.1 | | | 6.2 | 4.6 |
| | | RBD | 62.4 | 22.4 | | | 8.0 | 5.0 | | | 5.7 | 7.9 |
| | | SD1 | 4.8 | 1.7 | | | 2.5 | 0.6 | | | 1.1 | 1.7 |
| | | SD2 | 0.7 | 0.8 | | | 0.5 | 0.1 | | | 4.0 | 4.1 |
| | | S2 | 0.0 | 2.7 | | | 0.1 | 0.1 | | | 0.1 | 2.0 |
| | C | NTD | 4.3 | 0.9 | | | 2.2 | 1.4 | | | 2.5 | 4.3 |
| | | RBD | 0.7 | 0.3 | | | 0.5 | 0.4 | | | 2.4 | 6.7 |
| | | SD1 | 0.9 | 0.4 | | | 0.9 | 0.3 | | | 3.4 | 2.5 |
| | | SD2 | 2.9 | 0.3 | | | 1.3 | 0.4 | | | 1.5 | 4.4 |
| | | S2 | 0.3 | 2.8 | | | 0.1 | 1.0 | | | 0.2 | 0.8 |
| pH5.5-2 | A | NTD | 0.7 | 0.3 | | | | | | | 5.5 | 5.8 |
| | | RBD | 1.8 | 1.0 | | | | | | | 4.9 | 4.8 |
| | | SD1 | 1.8 | 0.1 | | | | | | | 12.0 | 3.4 |
| | | SD2 | 0.4 | 0.1 | | | | | | | 2.8 | 4.5 |
| | | S2 | 0.1 | 4.0 | | | | | | | 0.1 | 0.8 |
| | B-up | NTD | 1.5 | 0.5 | | | | | | | 6.3 | 4.6 |
| | | RBD | 64.6 | 25.0 | | | | | | | 2.9 | 7.1 |
| | | SD1 | 6.1 | 2.2 | | | | | | | 2.7 | 1.5 |
| | | SD2 | 0.8 | 0.7 | | | | | | | 3.6 | 4.0 |
| | | S2 | 0.2 | 2.8 | | | | | | | 0.2 | 2.0 |
| | C | NTD | 4.1 | 1.3 | | | | | | | 0.6 | 4.2 |
| | | RBD | 1.3 | 0.6 | | | | | | | 1.9 | 6.4 |
| | | SD1 | 0.9 | 0.7 | | | | | | | 2.6 | 2.2 |
| | | SD2 | 3.2 | 0.2 | | | | | | | 0.4 | 4.4 |
| | | S2 | 0.2 | 2.1 | | | | | | | 0.2 | 1.7 |

**Color coding:**

| ° or Å |
|---|
| < 1 |
| 1 - 3 |
| 3 - 5 |
| 5 - 20 |
| 20 - 100 |

* Movements are expressed as rotation angles and displacements between corresponding domains in the pH 5.5 and pH 4.0 structures. Structures were first superimposed in the S2 domain with residues 728 to 1140, domains within the same superimposed protomer were compared.

**Table S5. Inter- and Intra-Protomer Interface Areas for Switches, Related to Figures 5 and 7.**

| Buried surface area (Å$^2$) | Switch A | | | | Switch B | | | | Switch C | | | |
|---|---|---|---|---|---|---|---|---|---|---|---|---|
| | Inter-protomer on C | | Intra-protomer on A | | Inter-protomer on A | | Intra-protomer on B | | Inter-protomer on B | | Intra-protomer on C | |
| pH 4.0 | SD1 | 367 | NTD | 76 | SD1 | 351 | NTD | 81 | SD1 | 364 | NTD | 80 |
| | SD2 | 202 | S2 | 458 | SD2 | 165 | S2 | 466 | SD2 | 209 | S2 | 460 |
| | Asp614 | 90 | | | Asp614 | 72 | | | Asp614 | 85 | | |
| | Total | 569 | | 534 | | 516 | | 547 | | 573 | | 539 |
| pH5.5-up 1 | SD1 | 117 | NTD | 0 | SD1 | 152 | NTD | 3 | SD1 | 227 | NTD | 0 |
| | SD2 | 82 | S2 | 350 | SD2 | 297 | S2 | 534 | SD2 | 95 | S2 | 538 |
| | Asp614 | 19 | | | Asp614 | 74 | | | Asp614 | 45 | | |
| | NAG1309 | | | | NAG1309 | 53 | | | NAG1309 | 10 | | |
| | Total | 199 | | 350 | | 502 | | 537 | | 332 | | 538 |
| pH5.5-up2 | SD1 | 139 | NTD | 0 | SD1 | 146 | NTD | 10 | SD1 | 239 | NTD | 0 |
| | SD2 | 81 | S2 | 367 | SD2 | 223 | S2 | 547 | SD2 | 130 | S2 | 524 |
| | Asp614 | 23 | | | Asp614 | 70 | | | Asp614 | 56 | | |
| | NAG1309 | | | | NAG1309 | 36 | | | NAG1309 | | | |
| | Total | 220 | | 367 | | 404 | | 557 | | 369 | | 524 |
